# Stress-primed secretory autophagy drives extracellular BDNF maturation

**DOI:** 10.1101/2020.05.13.090514

**Authors:** Silvia Martinelli, Elmira A. Anderzhanova, Svenja Wiechmann, Frederik Dethloff, Katja Weckmann, Thomas Bajaj, Jakob Hartmann, Kathrin Hafner, Max L. Pöhlmann, Lee Jollans, Giuseppina Maccarrone, Felix Hausch, Christoph W Turck, Alexandra Philipsen, Mathias V. Schmidt, Bernhard Kuster, Nils C Gassen

## Abstract

The stress response is an essential mechanism that strives to maintain homeostasis, and its disruption is implicated in several psychiatric disorders. As a cellular response to stressors, autophagy is activated to regulate homeostasis through protein degradation and recycling. Secretory autophagy is a recently described pathway where autophagosomes fuse with the plasma membrane rather than lysosomes. In this study, we demonstrate that glucocorticoid-mediated stress enhances secretory autophagy, via the stress-responsive co-chaperone FK506-binding protein 51. We identified the matrix metalloproteinase 9 (MMP9) as one of the stress-induced secreted proteins. Using cellular assays and *in vivo* microdialysis, we further found that stress-enhanced MMP9 secretion increases the cleavage of pro-brain derived neurotrophic factor (proBDNF) to its mature form. BDNF is essential for adult synaptic plasticity and its pathway is associated with major depression and posttraumatic stress disorder. These findings unravel a novel mechanistic link between stress, stress adaptation and the development of psychiatric disorders, with possible therapeutic implications.

## Introduction

Excessive or prolonged stress represents a threat to homeostasis. To adapt to stress, several strategies evolved ranging from genetic and epigenetic changes to the activation of molecular pathways and the modification of physiological and social responses. Failure in stress adaptation can result in exaggerated stress responses that can lead to the development of numerous psychiatric disorders including major depressive disorder (MDD), anxiety, posttraumatic stress disorder (PTSD), bipolar disorder and schizophrenia^1^.

A fundamental molecular mechanism of stress adaptation is the maintenance of proteostasis, which is regulated by two major systems for protein turnover: 1) lysosomal degradation, such as autophagy, and 2) ubiquitin proteasomal degradation. By maintaining the balance between synthesis and degradation, these two processes contribute to the regulation of synaptic connectivity and plasticity in the central nervous system (CNS)^2,3^. Different types of lytic autophagy with different strategies for substrate selectivity preserve a healthy neural environment by preventing accumulation of protein aggregates, controlling the quality and quantity of cellular organelles, and complementing immune functions, such as pathogen defense^4^. Impairment of proteostatic regulation in the brain contributes to the development of proteinopathies and excessive inflammatory responses, hallmarks of neurodegenerative and neuropsychiatric diseases^5,6^. Cellular stressors including starvation, oxidative stress and infection are known threats to proteostasis that are counteracted by autophagy. We previously reported that, similarly to cellular stressors, glucocorticoid (GC)-mediated stress leads to the activation of macroautophagy, regulated by the stress-sensitive co-chaperone FK506-binding protein 51 (FKBP51, coded by *FKBP5*)^7^.

Recently, a novel, non-lytic type of autophagy, termed secretory autophagy, was described^8^. Starvation induces loss of lysosomal integrity, which derails the cargo proteins within autophagic vesicles to the plasma membrane for subsequent secretion into the extracellular milieu, instead of their degradation via lysosome-dependent lytic autophagy. Previous work has linked secretory autophagy to the immune response and inflammation^9^. Indeed, validated cargo proteins include potent inflammatory mediators such as cytokines and cathepsins^10–12^. Whether the main function of secretory autophagy is to discard the secreted proteins or relocate them to exert extracellular functions remains unclear^9–12^.

In this study, we investigated whether GC-mediated stress affects secretory autophagy, analogously to macroautophagy. Furthermore, we explored a possible role for secretory autophagy as a mechanism linking GC-mediated stress to the development of psychiatric disorders. Our results demonstrate that GC-mediated stress enhances secretory autophagy via FKBP51, and that the matrix metalloproteinase 9 (MMP9) is among the regulated proteins, which leads to an increase in cleavage of pro-brain-derived neurotrophic factor (proBDNF) to its mature form (mBDNF) both *in vitro* and *in vivo*. Hereby, we unveil a novel mechanism linking the GC-mediated stress response to neuroplasticity and cognitive function – key features intimately connected to stress adaptation and psychiatric disorders.

## Results

### FKBP51 mediates vesicle fusion underlying secretory autophagy

We previously identified the co-chaperone FKBP51 as an essential molecular link between GC-mediated stress and macroautophagy^7^. In order to gain more insight into the FKBP51-directed molecular environment, we sought to identify interaction partners of FKBP51 and defined its interactome. In HEK293 cells, we overexpressed FLAG-tagged FKBP51 followed by immunoprecipitation (IP) and mass spectrometry analysis of immunoprecipitates and identified 29 interactors (Supplementary Table S1). To determine possible links of these interactors to proteostasis-relevant pathways, we subjected the 29 interactors to an automated literature-mining screen for “autophagy”, “proteostasis”, and “ubiquitin proteasome system” using a custom-developed Python algorithm (Fig. 1a). Out of the proteins related to autophagy, the vesicular R-SNARE protein SEC22B piqued our interest due to its central function in the secretory autophagy pathway^13^. This release pathway involves a stepwise succession of signaling proteins, cargo receptors and RQ-SNARE fusion proteins for the secretion of vesicular cargo^13^. In short, the secretory autophagy cargo-receptor TRIM16 interacts with the R-SNARE SEC22B to recruit the cargo to LC3-II-positive sequestration membranes. Interaction of SEC22B with a Qabc-SNARE complex leads to vesicle-plasma membrane fusion, which further succeeds in cargo secretion. To investigate the role that FKBP51 plays in secretory autophagy, we first validated our interactome results via co-IP experiments in neuroblastoma SH-SY5Y cells. We confirmed, and validated for the first time, the interaction of FKBP51 with SEC22B (Fig. 1b,c), previously only deduced via differential centrifugation^13^, and with two additional FKBP51 interactors, RACK1 and UBC12 (Supplementary Fig. S1a). Insight into FKBP51’s impact on secretory autophagy regulation was gained by examining the interaction of FKBP51 with other major players of this pathway: the established secretory autophagy cargo cathepsin D (CTSD)^13^, its receptor TRIM16, and the autophagosomal membrane-spiking protein LC3B. Neither CTSD nor LC3B could be identified as an FKBP51 interactor in co-IP experiments (Fig. 1d), suggesting that FKBP51 does not contribute to the transport of cargo proteins or the decoration of autophagic vesicles by LC3B. TRIM16 was found to weakly bind to FKBP51, represented by a faint band, and confirmed in a reverse co-IP experiment (Supplementary Fig. S1b) prompting us to further investigate a putative role of FKBP51 in cargo-receptor dynamics. Thus, we generated an FKBP51-deficient SH-SY5Y cell line (Supplementary Fig. S1c) and analyzed the interaction of TRIM16 with CTSD and with SEC22B in the absence of FKBP51. Co-IP analyses revealed that FKBP51 does not affect the interactions of TRIM16 with its binding partners (Fig. 1e,f).

**Figure 1.**
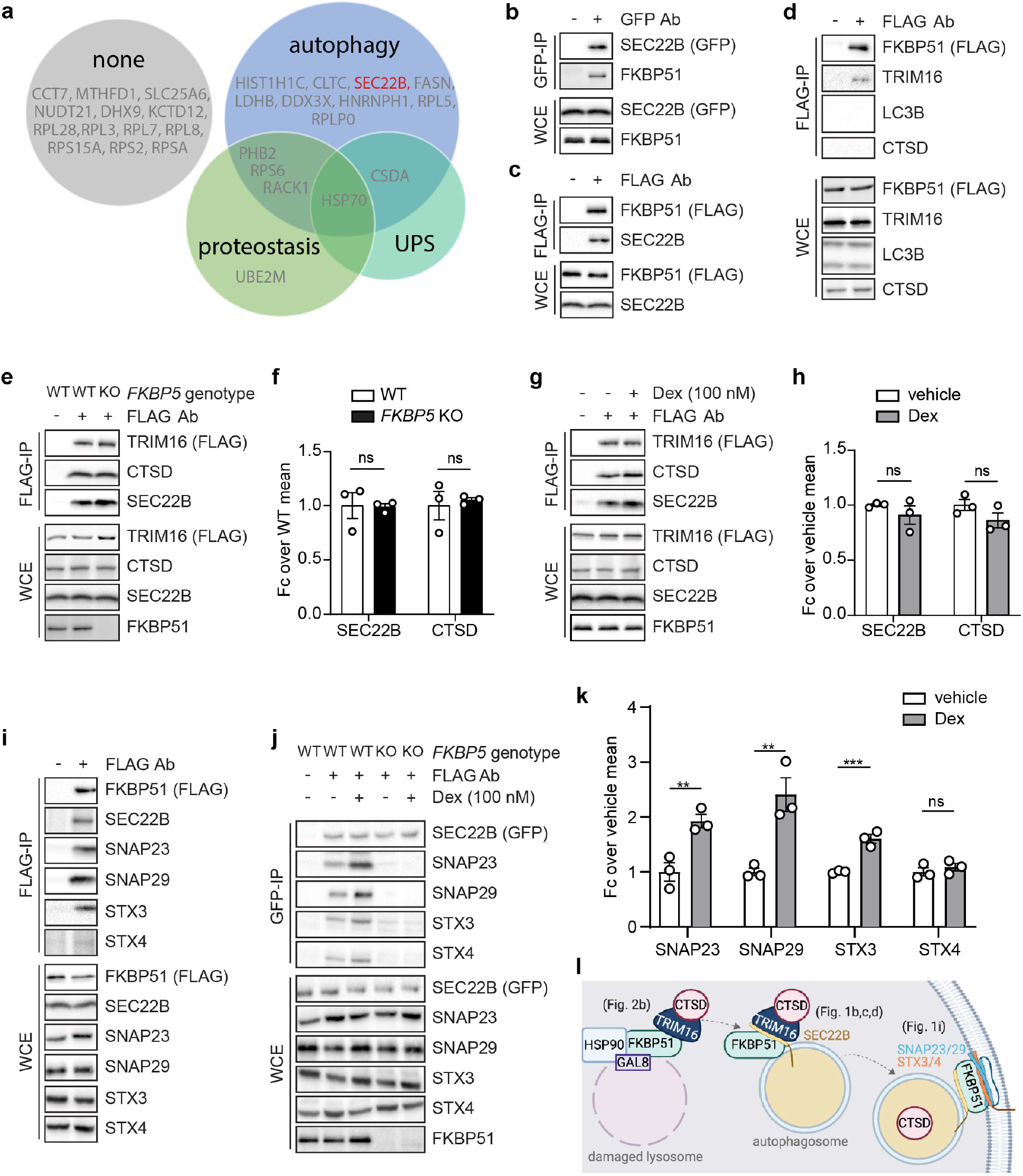
FKBP51 links stress to secretory autophagy. **a)** Results of automated literature mining of FKBP51 interactors in association to “autophagy”, “proteostasis” and “ubiquitin proteasome system”. **b)** Western blotting for FKBP51 and SEC22B in GFP-tagged SEC22B co-IP (GFP-IP) and whole cell extract (WCE) as control; **c)** Western blotting for FKBP51 and SEC22B in FLAG-tagged FKBP51 co-IP (FLAG-IP) and WCE as control. **d)** Western blotting for FKBP51, TRIM16, LC3B and CTSD in FLAG-tagged FKBP51 co-IP (FLAG-IP) and WCE as control. **e)** Western blotting for TRIM16, CTSD and SEC22B in FLAG-tagged TRIM16 co-IP (FLAG-IP) and WCE as control performed in WT and *FKBP5* KO cells. **f)** Quantifications of **e** with *n* = 3 per group. **g)** Western blotting for TRIM16, CTSD and SEC22B in FLAG-tagged TRIM16 co-IP (FLAG-IP) and WCE as control performed in cells treated with 100 nM dexamethasone or vehicle for 4 hours. **h)** Quantifications of **g** with *n* = 3 per group. **i)** Western blotting for FKBP51, SEC22B, SNAP23, SNAP29, STX3 and STX4 in FLAG-tagged FKBP51 co-IP (FLAG-IP) and WCE as control. **j)** Western blotting for, SEC22B, SNAP23, SNAP29, STX3 and STX4 in GFP-tagged SEC22B co-IP (GFP-IP) and WCE as control performed in WT and *FKBP5* KO cells treated with 100 nM dexamethasone or vehicle for 4 hours. **k)** Quantifications of **j** with *n* = 3 per group. **l)** Schematic overview of the interactions of FKBP51 in the secretory autophagy pathway. Unpaired t-tests were performed on all quantifications; ns= not significant, **P < 0.01, ***P < 0.001. Data shown as mean ± s.e.m. Ab, antibody; Fc, fold change.

Glucocorticoid receptor (GR) activation enhances FKBP51 expression, which mediates the effects of stress on different cellular pathways and functions. Here, we investigated whether GR activation influences the interactions between TRIM16 and CTSD or SEC22B by stimulating the cells with 100 nM dexamethasone or vehicle for four hours. Co-IP analyses showed that dexamethasone does not affect the interaction between TRIM16 and SEC22B nor between TRIM16 and CTSD (Fig. 1g,h).

The final step in secretory autophagy that allows for cargo secretion, is the SNARE complex formation between the R-SNARE SEC22B and the Q_abc_ SNARE complex, formed by the synaptosomal-associated proteins SNAP23 and SNAP29 and the syntaxins 3 and 4 (STX 3/4). This event leads to the fusion of the autophagosome with the plasma membrane, and subsequent release of the cargo proteins into the extracellular milieu. Since we identified SEC22B as a binding partner of FKBP51, we proceeded to examine the impact of FKBP51 on the RQ-SNARE fusion process. Via co-IP analyses, we detected strong interactions of FKBP51 with SNAP23, SNAP29 and STX3, and weaker binding to STX4 (Fig. 1i). Additionally, dexamethasone treatment strengthened RQ-SNARE interactions (Fig. 1j,k), presumably via FKBP51, which was enhanced upon dexamethasone treatment (Supplementary Fig. S1d). Interestingly, in the absence of FKBP51, the RQ-SNARE complex was almost abolished, both at basal condition and after dexamethasone induction, as shown by co-IP analyses of SEC22B and RQ-SNARE proteins in *FKBP5* KO cells (Fig. 1j,k). From these data, FKBP51 results to be involved in several key steps of the secretory autophagy pathway (Fig. 1l).

### GCs induce lysosomal damage triggering secretory autophagy

Another hallmark of secretory autophagy that plays a decisive role on whether autophagosomes undergo conventional lytic autophagy or fuse with cytoplasmic membranes for non-canonical secretion is the reduction of lysosomal integrity^13^. In order to test if GC-mediated endocrine stress affects lysosomal integrity, we analyzed in SH-SY5Y cells the effects of dexamethasone on the expression levels of galectin-3 (GAL3) and galectin-8 (GAL8), established markers of damaged endomembranes such as lysosomes^14,15^. Western blot analyses showed an overall increase in both GAL3 and GAL8 after dexamethasone treatment, suggesting enhanced lysosomal damage (Fig. 2a). Next, we determined whether FKBP51 interacts with GAL3 and/or GAL8. Via co-IP, we identified GAL8 as an FKBP51 interactor, whereas GAL3 did not appear to bind to FKBP51. To further specify the binding of FKBP51 to GAL8 we used a mutant FKBP51 lacking its ability to bind HSP90 (TPRmut). Co-IP of FKBP51/TPRmut and GAL8 revealed a decrease in GAL8 binding, suggesting that this interaction is (at least) partially mediated through HSP90 (Fig. 2b), which has been reported to co-localize with GAL8^13^.

**Figure 2.**
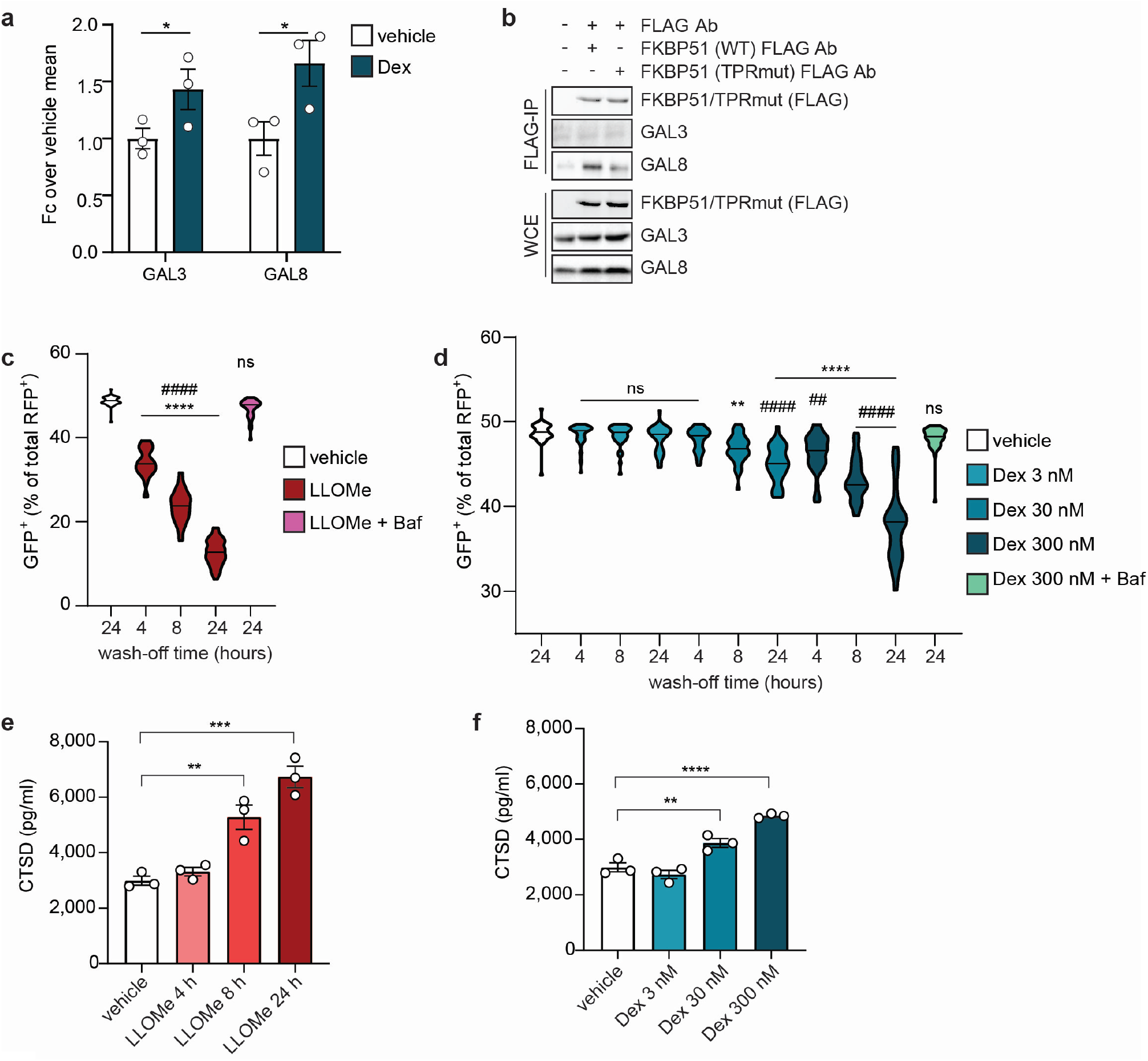
FKBP51 mediates regulatory mechanisms underlying secretory autophagy. **a)** Quantification of western blot analyses for GAL3 and GAL8 normalized to actin from SH-SY5Y WT cells treated with 100 nM dexamethasone (Dex) or vehicle for 4 hours. *n* = 3 per group. Unpaired t-tests were performed; *P < 0.05. Data shown as mean ± s.e.m. **b)** Western blotting for FKBP51, GAL3 and GAL8 in FLAG-tagged WT or mutant (TPRmut) FKBP51 co-IP (FLAG-IP) in SH-SY5Y and whole cell extract (WCE) as control performed in cells treated with 100 nM Dex or vehicle. **c)** and **d)** Quantification of GFP^+^ puncta expressed as a percentage of total RFP^+^ puncta in SH-SY5Y cells (*n* = 40 per group) transfected with tfGal3 construct and treated with 1 mM LLOMe or 300 nM Dex) for 3 hours, followed by 4, 8 and 24 hours wash-off, and with co-treatment of bafilomycin (Baf) for 3 hours followed by 24 hours wash-off. Kruskal-Wallis multiple comparison test was performed; **P < 0.01; ****P < 0.0001. * indicate comparisons to vehicle; # indicate comparisons to treatment + Baf. **e)** and **f)** CTSD from supernatants measured via ELISA after SIM-A9 cells were treated with LLOMe for 4, 8 and 24 hours or vehicle for 24 hours, or with 3 nM, 30 nM and 300 nM Dex or vehicle for 4 hours. *n* = 3 per group. Tukey’s multiple comparison test was performed; **P < 0.01; ***P < 0.001; ****P < 0.0001. * indicate comparisons to vehicle; # indicate comparisons to treatment + Baf. Data shown as mean ± s.e.m. Fc, fold change; Ab, antibody.

To directly assess lysosomal damage, we used tfGal3 (tandem fluorescent-tagged Galectin3), an established reporter system using an mRFP- and EGFP-tagged GAL3 in SH-SY5Y cells. Since mRFP and EGFP display differential sensitivity to acidic environments, tfGal3 enables the monitoring of pH change in compromised lysosomes^16^. After transient transfection with tfGal3, various treatments were applied for four hours and subsequently washed off at different time periods as indicated in Fig. 2c and d. L-leucyl-L-leucine methyl ester (LLOMe) and bafilomycin A1 (Baf) were used for assay calibration^16^. LLOMe is a lysosome-damaging and inflammasome-activating substance, while Baf inhibits the vacuolar V-ATPase, thereby preventing lysosomal acidification^17^. Using fluorescence microscopy, EGFP^+^ and mRFP^+^ puncta were quantified and set in ratio. A reduction in EGFP^+^ puncta was detected upon LLOMe treatment, indicating severe loss of lysosomal integrity that could be rescued through co-application of Baf (Fig. 2c).

Interestingly, dexamethasone caused a similar decrease in EGFP^+^ puncta with increasing concentrations (30 nM and 300 nM) and extended time after wash-off (8 and 24 hours), indicating induced lysosomal damage (Fig. 2d). This effect could be rescued by Baf (Fig. 2d). The obtained results indicate that GR activation by GCs induces lysosomal damage. To determine the effect of GC-induced lysosomal damage on secretory autophagy, we analyzed CTSD release from cells using enzyme-linked immunosorbent assay (ELISA). Murine microglia cell line SIM-A9 was employed for its high secretory capacity, and confirmed the interaction between FKBP51, SEC22B and the Q-SNARE proteins (Supplementary Fig. S2). In line with our previous results, the same treatments that caused an increase in lysosomal damage, also led to enhanced secretion of CTSD (Fig. 2e,f), suggesting that GR activation triggers secretory autophagy.

### Identification of novel cargo proteins regulated by stress-induced secretory autophagy

In order to capture the whole range of GC-mediated stress effects on secretory autophagy, and to identify the entire spectrum of cargo proteins released by this mechanism, we performed a secretome-wide analysis. We controlled for autophagy-dependent secretion of the analyzed proteins using a CRISPR-Cas9-generated autophagy-deficient microglial cell line (*Atg5* KO SIM-A9; Supplementary Fig. S3). This cellular model was further verified by quantification of secreted CTSD levels via ELISA. We observed enhanced secretion of CTSD in response to 100 nM dexamethasone in WT cells, while this increase was annulled in the absence of ATG5 (Fig. 3a), demonstrating that dexamethasone-induced secretory autophagy is tightly linked to ATG5-mediated signaling, and confirming the validity of this model.

**Figure 3.**
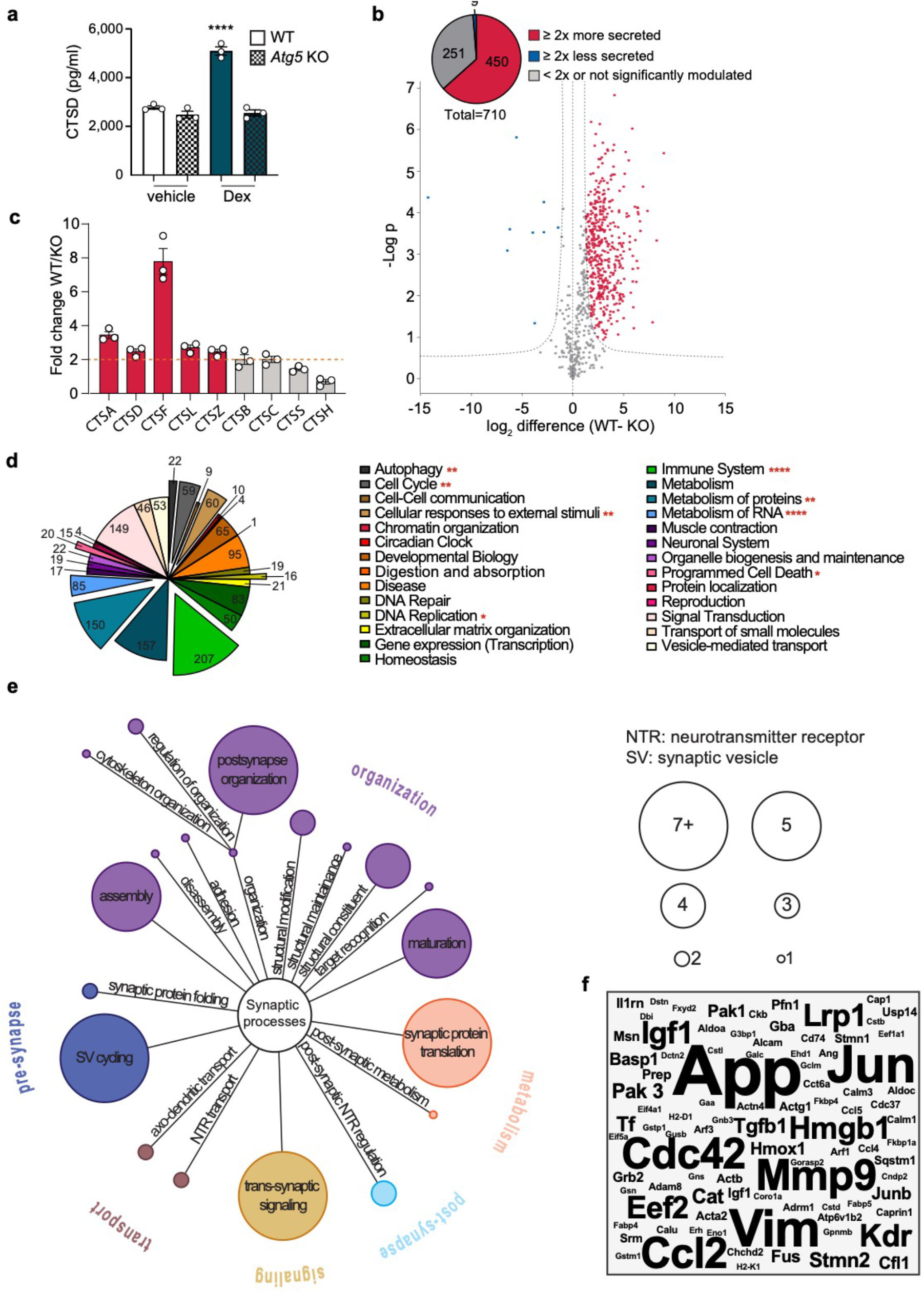
Detection of novel cargo proteins regulated by stress-induced secretory autophagy. **a)** CTSD from supernatants measured via ELISA after WT or *Atg5* KO SIM-A9 cells were treated with 300 nM dexamethasone (Dex) or vehicle for 4 hours. Tukey’s multiple comparison test was performed; *P < 0.05. Data shown as mean ± s.e.m. **b)** Volcano plot representation of a multiple t-test analysis of the secretome (difference of WT-*Atg5* KO); FDR=0.01, s0=1. **c)** Secretion levels of detected cathepsins indicated as fold change of WT over *Atg5* KO. **d)** GO results performed with Reactome. **e)** GO results performed with SynGO. **f)** Word cloud representation of automated literature mining results in association with “neuroplasticity”.

For the secretome analysis, we used a proteomic method that combines metabolic labeling and click chemistry to selectively enrich and quantify released proteins^18^. Secretory autophagy was induced by treatment of WT and *Atg5* KO cells with dexamethasone. After LC-MS/MS measurement, database search and quality filtering, we were able to identify a total of 710 secreted proteins. A multiple t-test analysis (Fig. 3b; Supplementary Table S2) revealed that the secretion of a large number of proteins (75% of the total = 530 proteins) was enhanced in the WT sample compared to the *Atg5* KO upon dexamethasone treatment (FDR<0.01). From this group, the vast majority (85% = 450 proteins) showed a change of at least 2-fold, while less than 4% of the detected proteins presented lower secretion levels in the WT condition compared to *Atg5* KO. Interestingly, we could not only confirm an increase in secretion of CTSD, but also identified several other cathepsins that presented a strongly enhanced autophagy-dependent secretion (Fig. 3c), thereby uncovering secretory autophagy as a highly relevant release mechanism for members of the cathepsin protein family from microglial cells. Among the nine detected cathepsins, only cathepsin H (CTSH; also identified as pro-CTSH) displayed a lower secretion from the WT cells as compared to *Atg5* KO cells (Fig. 3c). For a better understanding of the functional profile of the autophagy-regulated secretome, we performed a gene ontology (GO) analysis of the 450 proteins with increased secretion. The results displayed in Figure 3d reveal a high functional heterogeneity of the analyzed set of proteins, covering most annotated pathways (full list in Supplementary Table S3). To pinpoint the role of GC-induced secretory autophagy in mechanisms of synaptic plasticity, we performed a synapse-specific GO analysis using the recently established SynGO analysis tool^19^ (Fig. 3e; full results in Supplementary Table S4). 72 proteins were matched to SynGO annotated data sets, covering an interestingly high number of synapse-relevant proteins. Almost half of these proteins (33 of 72) were found to be annotated in the cluster of synaptic organization, playing an essential role in structural reorganization and neurite growth, key features of synaptic plasticity. To further verify these data and identify neuroplasticity-related proteins linked to stress-induced secretory autophagy, we performed an automated literature screening of the 450 analyzed proteins in association with neuroplasticity. Strikingly, 161 proteins were associated to neuroplasticity by at least one peer-reviewed publication (Fig. 3f; full results in Supplementary Table S5).

For a better understanding of the resulting neuroplasticity-related proteins, we manually analyzed and summarized the functions of the proteins with at least five publications (Table 1). Vimentin and MMP9 were of particular interest due to their reported extracellular function. Of notice, MMP9 is an important regulator in BDNF signaling^20^ and a candidate factor of stress-related changes in the brain^21^. Furthermore, over 70% of the top-ranked proteins (highlighted in green) were also linked to the immune system (as resulted by the previous GO analysis). This result strengthened the evidence linking secretory autophagy to the immune response and further broadened their association to neuroplasticity.

**Table 1.**
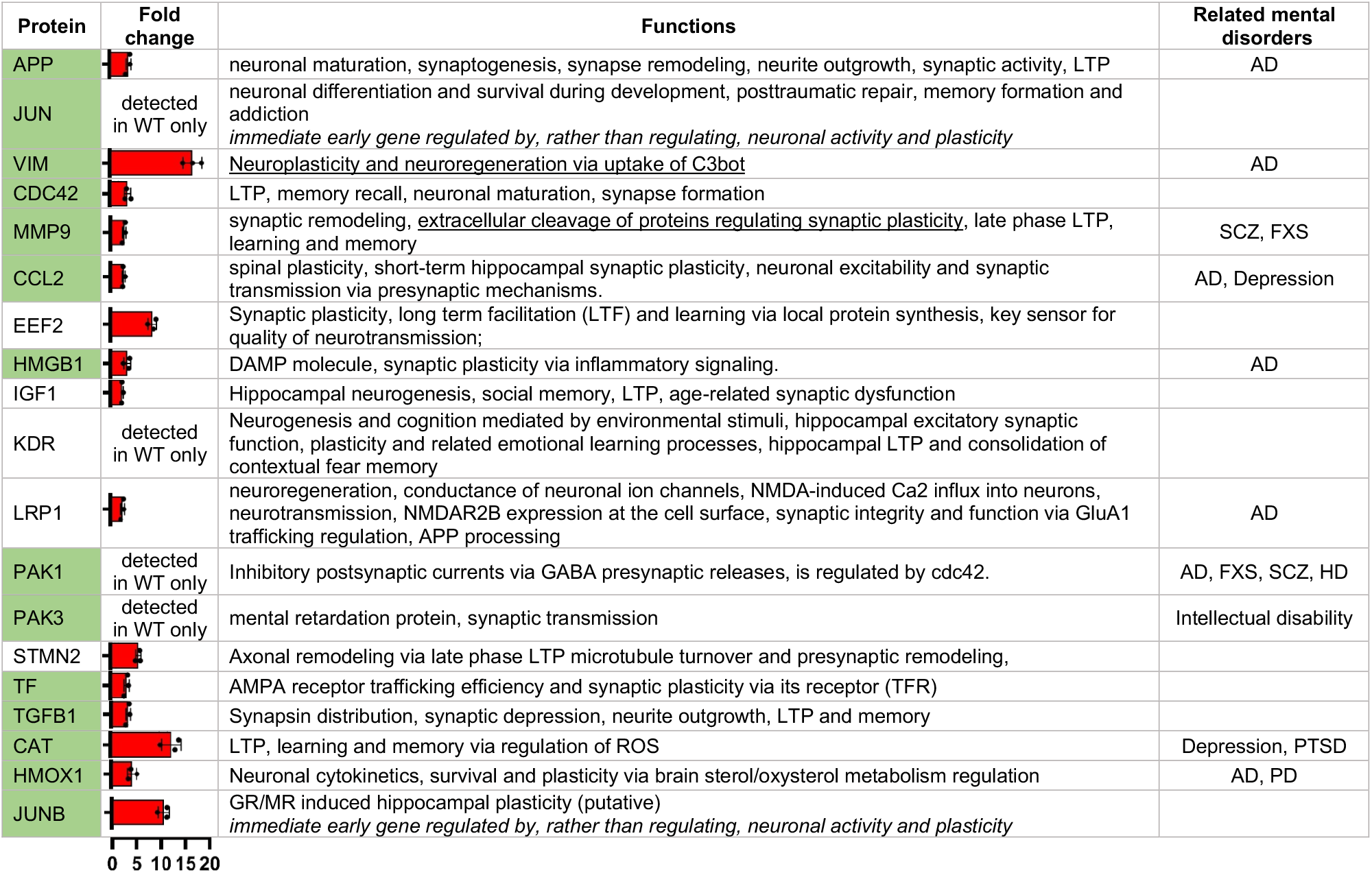
Neuroplasticity-related proteins regulated by stress-induced secretory autophagy. List of proteins and related functions found to be associated to neuroplasticity in at least five peer-reviewed publications. Bars indicate fold change in secretion of WT over Atg5 KO cells. Green background indicates proteins also involved in immune response based on GO analysis; underlining indicates extracellular function. AD, Alzheimer’s disease; PD, Parkinson’s disease; SCZ, schizophrenia; HD, Huntington disease; PTSD, Post traumatic stress disorder; FXS, fragile X syndrome.

| Protein | Fold change | Functions | Related mental disorders |
| --- | --- | --- | --- |
| APP |  | neuronal maturation, synaptogenesis, synapse remodeling, neurite outgrowth, synaptic activity, LTP | AD |
| JUN | detected in WT only | neuronal differentiation and survival during development, posttraumatic repair, memory formation and addiction<br>*immediate early gene regulated by, rather than regulating, neuronal activity and plasticity* |  |
| VIM |  | <u>Neuroplasticity and neuroregeneration via uptake of C3bot</u> | AD |
| CDC42 |  | LTP, memory recall, neuronal maturation, synapse formation |  |
| MMP9 |  | synaptic remodeling, <u>extracellular cleavage of proteins regulating synaptic plasticity</u>, late phase LTP, learning and memory | SCZ, FXS |
| CCL2 |  | spinal plasticity, short-term hippocampal synaptic plasticity, neuronal excitability and synaptic transmission via presynaptic mechanisms. | AD, Depression |
| EEF2 |  | Synaptic plasticity, long term facilitation (LTF) and learning via local protein synthesis, key sensor for quality of neurotransmission; |  |
| HMGB1 |  | DAMP molecule, synaptic plasticity via inflammatory signaling. | AD |
| IGF1 |  | Hippocampal neurogenesis, social memory, LTP, age-related synaptic dysfunction |  |
| KDR | detected in WT only | Neurogenesis and cognition mediated by environmental stimuli, hippocampal excitatory synaptic function, plasticity and related emotional learning processes, hippocampal LTP and consolidation of contextual fear memory |  |
| LRP1 |  | neuroregeneration, conductance of neuronal ion channels, NMDA-induced Ca2 influx into neurons, neurotransmission, NMDAR2B expression at the cell surface, synaptic integrity and function via GluA1 trafficking regulation, APP processing | AD |
| PAK1 | detected in WT only | Inhibitory postsynaptic currents via GABA presynaptic releases, is regulated by cdc42. | AD, FXS, SCZ, HD |
| PAK3 | detected in WT only | mental retardation protein, synaptic transmission | Intellectual disability |
| STMN2 |  | Axonal remodeling via late phase LTP microtubule turnover and presynaptic remodeling, |  |
| TF |  | AMPA receptor trafficking efficiency and synaptic plasticity via its receptor (TFR) |  |
| TGFB1 |  | Synapsin distribution, synaptic depression, neurite outgrowth, LTP and memory |  |
| CAT |  | LTP, learning and memory via regulation of ROS | Depression, PTSD |
| HMOX1 |  | Neuronal cytokinetics, survival and plasticity via brain sterol/oxysterol metabolism regulation | AD, PD |
| JUNB |  | GR/MR induced hippocampal plasticity (putative)<br>*immediate early gene regulated by, rather than regulating, neuronal activity and plasticity* |  |

### Stress enhances mBDNF production by promoting proBDNF cleavage via extracellular MMP9

The above findings led us to hypothesize a model whereby stress-induced release of MMP9 through secretory autophagy increases cleavage of proBDNF to mBDNF. To validate this hypothesis, we first determined the levels of proBDNF and mBDNF as well as MMP9 in the supernatants of WT and *Atg5* KO SIM-A9 cells exposed to dexamethasone. Both, MMP9 and mBDNF were significantly more abundant in the supernatants of WT cells upon dexamethasone stimulation, but not in the supernatants of *Atg5* KO cells, as shown by ELISA (Fig. 4a,c). Conversely, proBDNF levels were slightly increased in response to dexamethasone, however in a genotype-independent manner (Fig. 4b). To exclude secondary effects of GCs (dexamethasone) on the secretion of BDNF and MMP9, in a second set of experiments we compared the effect of the more selective lysosomotropic agent LLOMe and tested different concentrations of dexamethasone (Supplementary Fig. S4a,b). These results confirmed that enhanced MMP9 levels correlate with increased levels of mBDNF, but not proBDNF. In order to verify that this increase is determined by enhanced cleavage of proBDNF to mBDNF, rather than increased secretion, we repeated the experiments in the presence of MMP9 inhibitor I (MMP9i). As expected, MMP9 secretion increased after dexamethasone treatment and was not affected by the presence of MMP9i (Fig. 4d), while proBDNF levels were unaffected both by the inhibition of MMP9 and by dexamethasone treatment (Fig. 4e). Conversely, while mBDNF levels increased after dexamethasone treatment (compared to vehicle), they were reduced upon co-application of MMP9i (Fig. 4f). Together these results strongly support our model, proposing that stress (dexamethasone) enhances MMP9 secretion leading to the conversion of proBDNF to mBDNF, thereby raising the mBDNF/proBDNF ratio by more than 2-fold compared to vehicle (Supplementary Fig.S4c).

**Figure 4.**
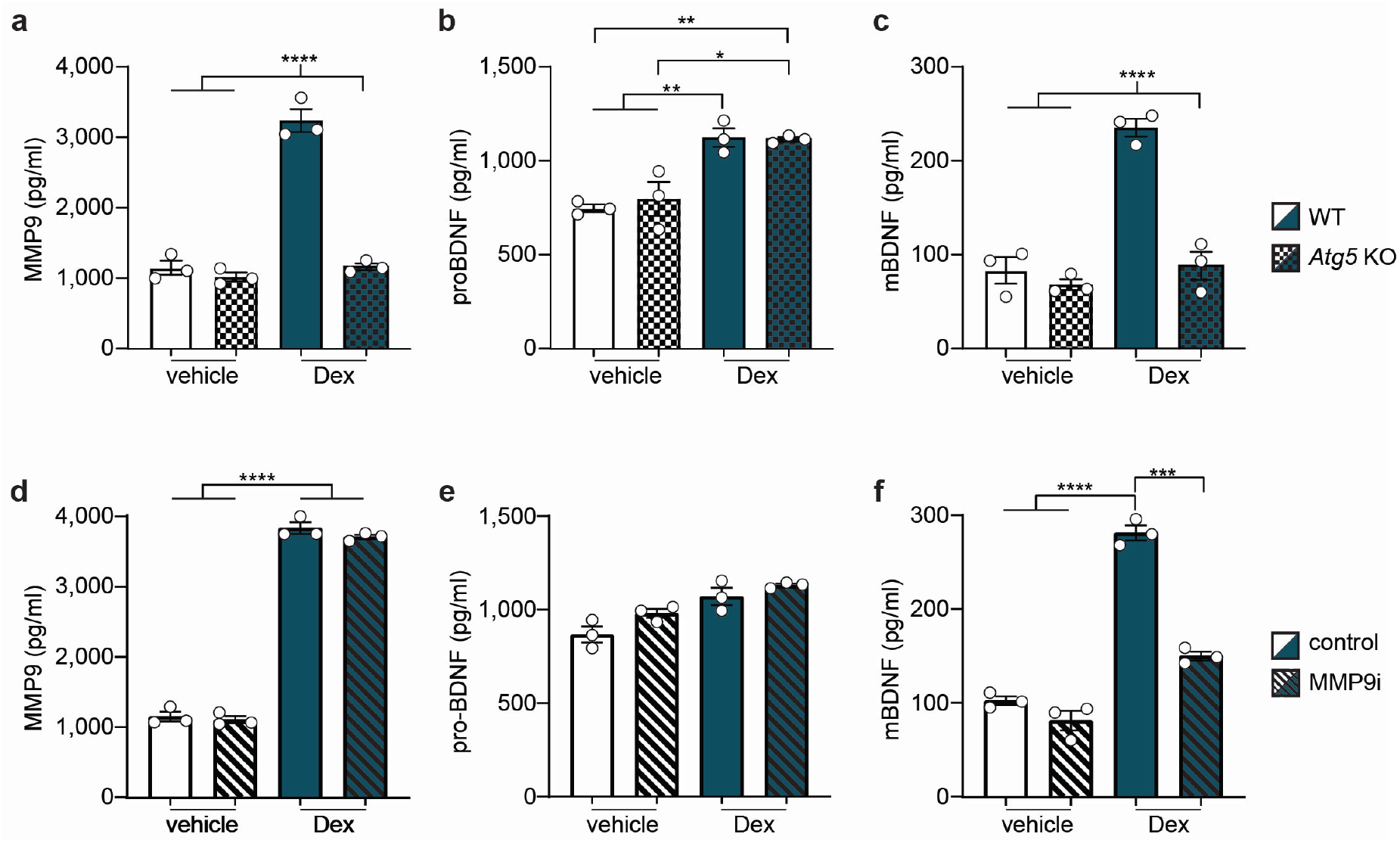
Stress enhances mBDNF production promoting proBDNF cleavage via MMP9. **a)** MMP9, **b)** proBDNF, **c)** mBDNF levels from supernatants measured via ELISA after WT or *Atg5* KO SIM-A9 cells were treated with 300 nM dexamethasone (Dex) or vehicle for 4 hours. **d)** MMP9, **e)** proBDNF, **f)** mBDNF levels from supernatants measured via ELISA after WT SIM-A9 cells were treated with 300 nM Dex, Dex + MMP9 inhibitor I (MMP9i), or vehicle for 4 hours. Tukey’s multiple comparison tests were performed; *P < 0.05; **P < 0.01; ***P < 0.001; ****P < 0.0001. Data shown as mean ± s.e.m.

### Stress enhances secretory autophagy and elevates extracellular mBDNF *in vivo*

To understand the relevance of stress-induced secretory autophagy and its impact on brain physiology, we used microdialysis, a powerful neurochemistry approach for monitoring changes in extracellular content of endogenous or exogenous substances in animals *in vivo* (Fig. 5a). With this translation to a murine model, we aimed not only to achieve a better understanding of the secretion dynamics of selected proteins in a complex system, but also to validate the molecular mechanism we discovered *in vitro*, by application of a physiological stressor in living animals. We exposed mice to an acute electric foot-shock (FS; 1.5 mA, × 2) and quantitatively analyzed changes in selected proteins in the extracellular space of the medial prefrontal cortex (mPFC). To assess the effect of acute stress on protein secretion dynamics, microdialysate fractions were collected during baseline condition (at 120, 60 and 30 minutes prior to the FS), at the onset of the FS and after the FS (post-FS; at 30 and 150 minutes post FS) (Fig. 5b). To control for the dependency of the analyzed secretion on the FKBP51-regulated secretory autophagy pathway characterized *in vitro*, we compared secretion dynamics of WT and *Fkbp5* KO mice. The 30-minute dialysate fractions were analyzed using capillary electrophoresis-based immunodetection (Wes, ProteinSimple). To verify this approach, we first analyzed the established secretory autophagy cargo CTSD. The results revealed increased CTSD levels after FS (Fig. 5c). This increase was strongly diminished in the *Fkbp5* KO animals, confirming our *in vitro* data that stress-induced secretion of CTSD is mediated by FKBP51. Furthermore, to confirm our *in vitro* data on the regulation of BDNF maturation *in vivo*, we analyzed MMP9, proBDNF and mBDNF in the microdialysates.

**Figure 5.**
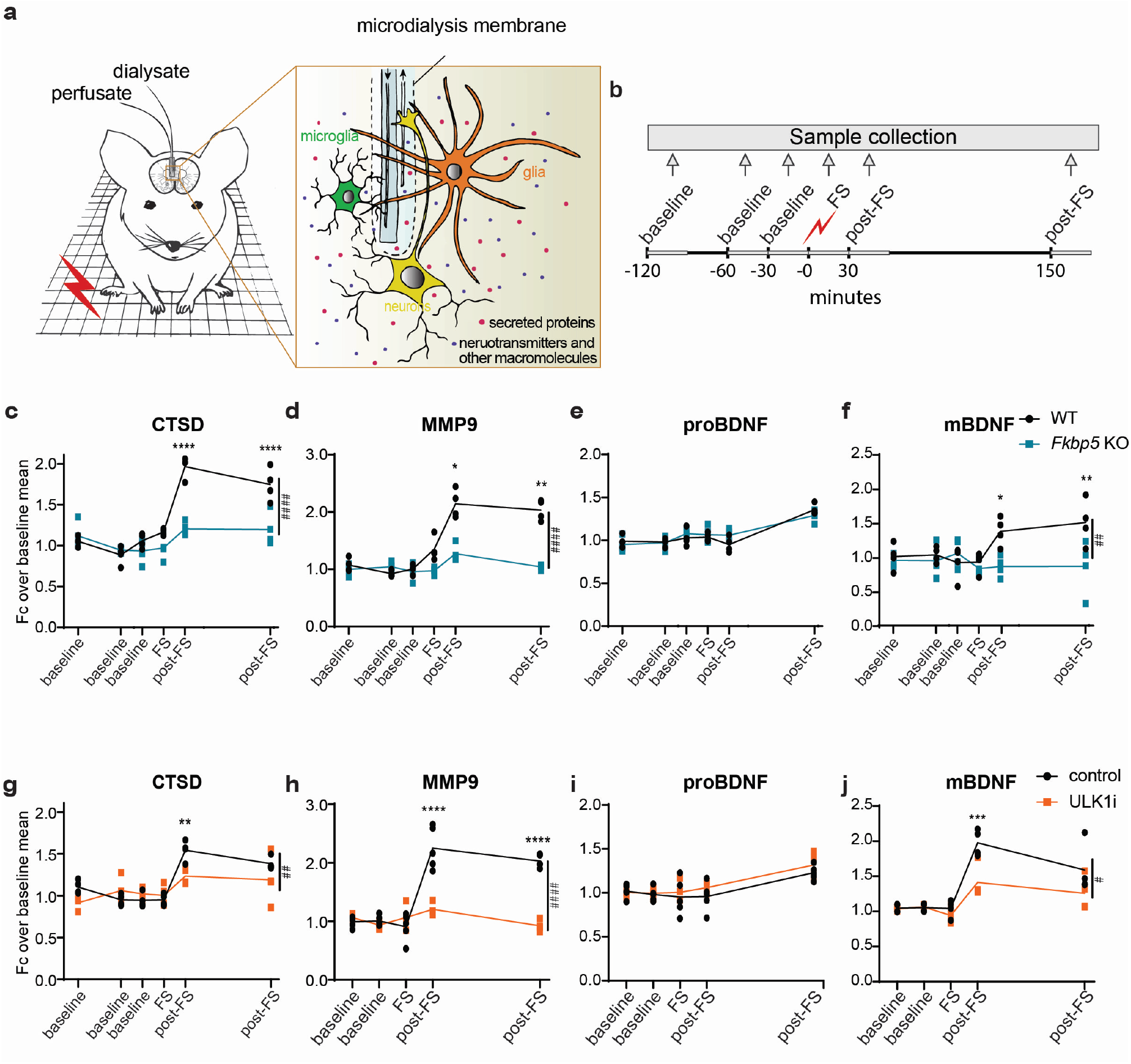
Stress enhances secretory autophagy and increases the mBDNF/proBDNF ratio *in vivo*. **a)** Schematic overview of *in vivo* microdialysis. **b)** Experimental design and; each sample was collected over 30 min indicated by the light grey lines. Quantifications of **c)** CTSD**, d)** MMP9, **e)** proBDNF, and **f)** mBDNF from *in vivo* mPFC microdialyses of WT and *Fkbp5* KO mice. Quantifications of **g)** CTSD, **h)** MMP9, **i)** proBDNF and **j)** mBDNF from *in vivo* mPFC microdialyses of WT mice injected intraperetoneally with ULK1 inhibitor (ULK1i) or saline. *n* = 4 mice per group.Two-way ANOVA with multiple t-test was performed; *P < 0.05; **P < 0.01; ***P < 0.001; ****P<0.0001. # refers to time x genotype/treatment interaction. Data shown as mean ± s.e.m. FS, foot shock; Fc, fold change.

Quantification of these proteins showed an increase of MMP9 and mBDNF levels after FS (Fig. 5d,f). In line with the results for CTSD, this increase was strongly diminished in the *Fkbp5* KO animals. ProBDNF showed a very late mild increase 150 min after FS, independently of the genotype (Fig. 5f). This phenomenon firstly confirmed that the modulation is specific for mBDNF, and therefore mediated by MMP9 (as also demonstrated *in vitro* via MMP9 inhibition; Fig. 4d,e,f), and secondly revealed an interesting regulation of proBDNF by stress independently of secretory autophagy, which we also observed *in vitro* (Supplementary Fig. S4b,S5). To confirm that secretion of CTSD and MMP9 and the respective changes in the extracellular mBDNF levels are dependent on the autophagic machinery, we measured their stress-dependent extracellular dynamics in the mPFC microdialysates of mice injected with the selective ULK1 inhibitor MRT 68921 (ULKi, 5 mg kg^−1^ intraperitoneally), an established inhibitor of autophagy^22^. An increase of CTSD and MMP9 was determined in the vehicle injected mice after FS (Fig. 5g,h). This induction was significantly impaired in mice treated with ULK1i. Accordingly, mBDNF followed an autophagy-dependent increase after FS (Fig. 5j), which corresponded to increased MMP9, while proBDNF showed a late-phase increase both in vehicle and ULK1i-treated mice (Fig. 5i). These data corroborate the *in vitro* findings showing that psychophysiological stress not only has a strong effect on autophagy-dependent secretion of CTSD and MMP9, but also leads to an increased extracellular mBDNF enrichment in murine mPFC.

### FKBP51 as target for stress-induced neuroplasticity modulation

Given that FKBP51 plays a central role in the regulation and mediation of stress on secretory autophagy, we tested pharmacological antagonization of this protein. We first verified the sole effect of increased FKBP51 on CTSD, MMP9, and BDNF secretion. We overexpressed FKBP51 in SIM-A9 cells and analyzed the supernatants with ELISA. Interestingly, higher levels of FKBP51 (mimicking the effect of GR activation by stress) were sufficient to cause enhanced secretion of CTSD and MMP9 (Fig. 6a,b). As with GCs, levels of mBDNF but not proBDNF were increased in the presence of high levels of FKBP51, indicating an elevated MMP9-dependent cleavage rather than increased secretion of the neurotrophin itself (Fig. 6c,d). Having confirmed the key function of FKBP51 in stress-mediated secretory autophagy, we tested the effect of the FKBP51 antagonist SAFit1^23^ on this mechanism. SIM-A9 cells were co-treated with SAFit1 or vehicle and dexamethasone or vehicle for four hours. ELISA quantifications indicated that SAFit1 treatments prevented the dexamethasone-induced CTSD and MMP9 secretion (Fig. 6e,f). Furthermore, SAFit1 significantly reduced the effect of dexamethasone on mBDNF but not on proBDNF (Fig. 6g,h). These results revealed FKBP51 as a potential drug target regulating BDNF-dependent neuroplasticity underlying stress-related psychiatric disorders.

**Figure 6.**
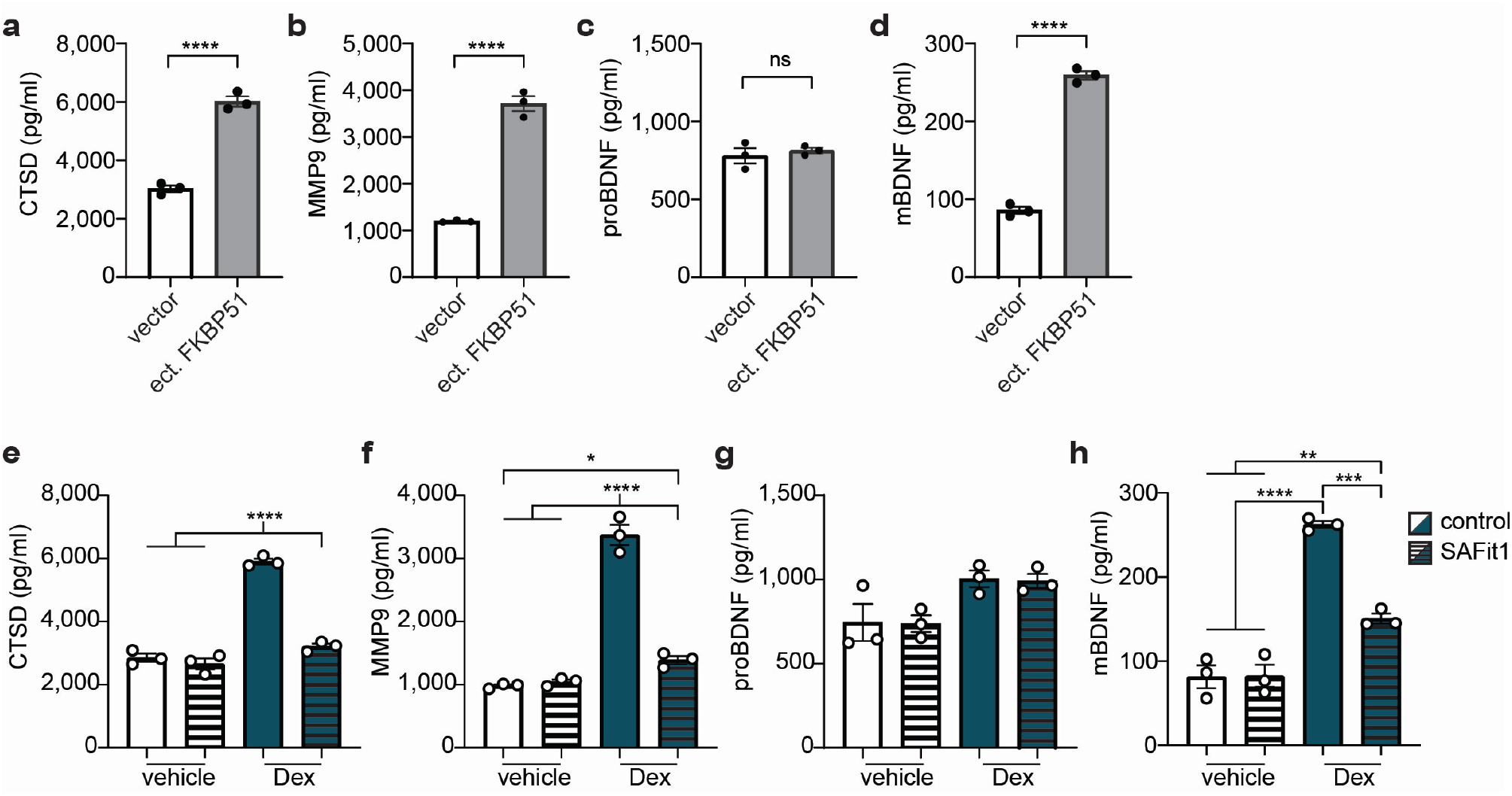
Targeting FKBP51 as possible treatment for stress-related pathologies. **a)** CTSD, **b)** MMP9, **c)** proBDNF and **d)** mBDNF levels from supernatants measured via ELISA after WT SIM-A9 cells were transfected with FKBP51 expressing plasmid (ect. FKBP51) or control vector. Unpaired t-tests were performed; ns= not significant, ****P < 0.0001; Data shown as mean ± s.e.m. **e)** CTSD, **f)** MMP9, **g)** proBDNF, **h)** mBDNF levels from supernatants measured via ELISA after WT SIM-A9 cells were treated with vehicle, SAFit, 300 nM dexamethasone (Dex) and 300 nM Dex + SAFit for 4 hours. Tukey’s multiple comparison test was performed; *P < 0.05; **P < 0.01; ***P < 0.001; ****P < 0.0001. Data shown as mean ± s.e.m.

## Discussion

In this study we describe a novel mechanism in which GC signaling and acute FS stress enhance secretory autophagy in an FKBP51-dependent manner, leading to increased extracellular BDNF maturation via elevated secretion of MMP9. BDNF is essential for synaptic plasticity processes which are required for long-term learning and memory, and stress coping. Notably, altered BDNF signaling has repeatedly been associated with stress-related psychopathology. Our findings may therefore be helpful to identify novel prevention and treatment approaches for psychiatric disorders including MDD and PTSD.

### Stress triggers secretory autophagy via FKBP51

Secretory autophagy is defined by three regulatory stages: 1) first, the disruption of lysosomal membranes by a stressor and the recruitment of TRIM16 and its cargo by galectins; 2) the transport of the TRIM16-cargo complex to the autophagosomal membrane via SEC22B; 3) the internalization of the cargo protein into the autophagosome, followed by its fusion with the plasma membrane mediated by the RQ-SNARE complex. This ultimately leads to the secretion of cargo proteins into the extracellular milieu. We identified the stress-responsive protein FKBP51 as a scaffolder and key driver of secretory autophagy throughout all its stages (Fig. 1l).

The first step, at the crossroads between macroautophagy and secretory autophagy, defines the cargo’s fate. Our previous study showed that stress enhances macroautophagy via FKBP51^7^. Here we find that high GC (dexamethasone) levels lead to increased GAL3/8 expression, which is indicative for increased lysosomal damage. GAL3 and GAL8 act as danger receptors and recruit TRIM16 in case of lysosomal damage^15,24^. This is in line with experiments using tfGal3 to monitor damaged lysosomes, which confirmed that dexamethasone stimulation indeed leads to reduced lysosomal integrity. These results indicate that sustained stress, as it occurs in disease progression of various psychiatric disorders such as PTSD, may lead to damage of lysosomal membranes, which in turn causes a switch from macroautophagy to secretory autophagy.

Co-IP experiments indicated that GAL8 interacts with FKBP51 suggesting that FKBP51 is recruited to damaged lysosomes and bridges the fusion to autophagosomes via SEC22B. In fact, we show that increased levels of lysosomal damage are paralleled by enhanced secretion of the established cargo CTSD. Our mechanistic analyses of the dexamethasone-triggered activation of secretory autophagy confirmed that FKBP51 is a key component and essential driver of the last step of secretory autophagy, by chaperoning the hetero-complex assembly of the vesicular R-SNARE SEC22B with the membranous Q-SNAREs enabling vesicle-membrane fusion. Furthermore, we observed a stronger and FKBP51-dependent interaction of the RQ-SNARE complex and an enhanced secretion of CTSD upon dexamethasone treatment. Taken together, these results confirm that GC-mediated stress enhances secretory autophagy via FKBP51.

The analysis of the GC-induced secretome revealed that a surprisingly high proportion of secreted proteins (63%) show an autophagy-dependent release, indicated by higher level of secretion in WT (2-fold or more) compared to *Atg5* KO. The subsequent GO analysis on this group of induced secreted proteins (450) showed a high functional heterogeneity, suggesting that secretory autophagy is not specific for the regulation of a particular functional protein group. Enrichments (FDR<0.05) in pathways of autophagy, cellular response to external stimuli and metabolism of proteins were expected given the high association of such functions with secretory autophagy and stress. Additionally, proteins related to the immune system were found to be enriched (FDR=1.91E-8), which is in line with previous evidence showing the role of secretory autophagy in extracellular signaling of the immune response^9,11,12,25^. Proteins involved in cell cycle, DNA replication, metabolism of RNA and programmed cell death were also enriched, which could reflect the cellular response and adaption to stress. The successive neuron-specific analysis performed with SynGO, revealed that 72 proteins have synaptic functions, of which almost half (33) is involved in synaptic organization. This result supported our hypothesis of stress-induced secretory autophagy having an effect on synaptic reorganization and neuroplasticity. Additionally, the literature mining resulted in 163 proteins with at least one peer-reviewed publication related to neuroplasticity. A deeper analysis of the top candidates further illustrated that a large number of the neuroplasticity-related proteins is associated with psychiatric and neurodegenerative disorders (Table 1). This strengthened our model for which stress might affect proteostasis underlying not only neurodegenerative disorders, but also psychiatric diseases, as proposed recently by Bradshaw and Korth^26^.

Based on our results and previous publications, we propose a dual-step-stress model in which an acute stress activates a first defense line that triggers lytic autophagy that in turn guarantees macromolecule recycling and degradation of potentially damaging aggregates. However, in the presence of prolonged or excessive stress, the cell switches to a second line of defense, which causes lysosomal damage and conveys the disposal of potentially damaging proteins to a secretory pathway, i.e., secretory autophagy. Furthermore, the finding that a large proportion of regulated proteins is involved in the immune response, suggests that an initially positive effect that increases neuroplasticity and cognitive arousal, would come with the cost of neuroinflammation in the case of prolonged or chronic stress (see schematic representation in Fig. 7).

**Figure 7.**
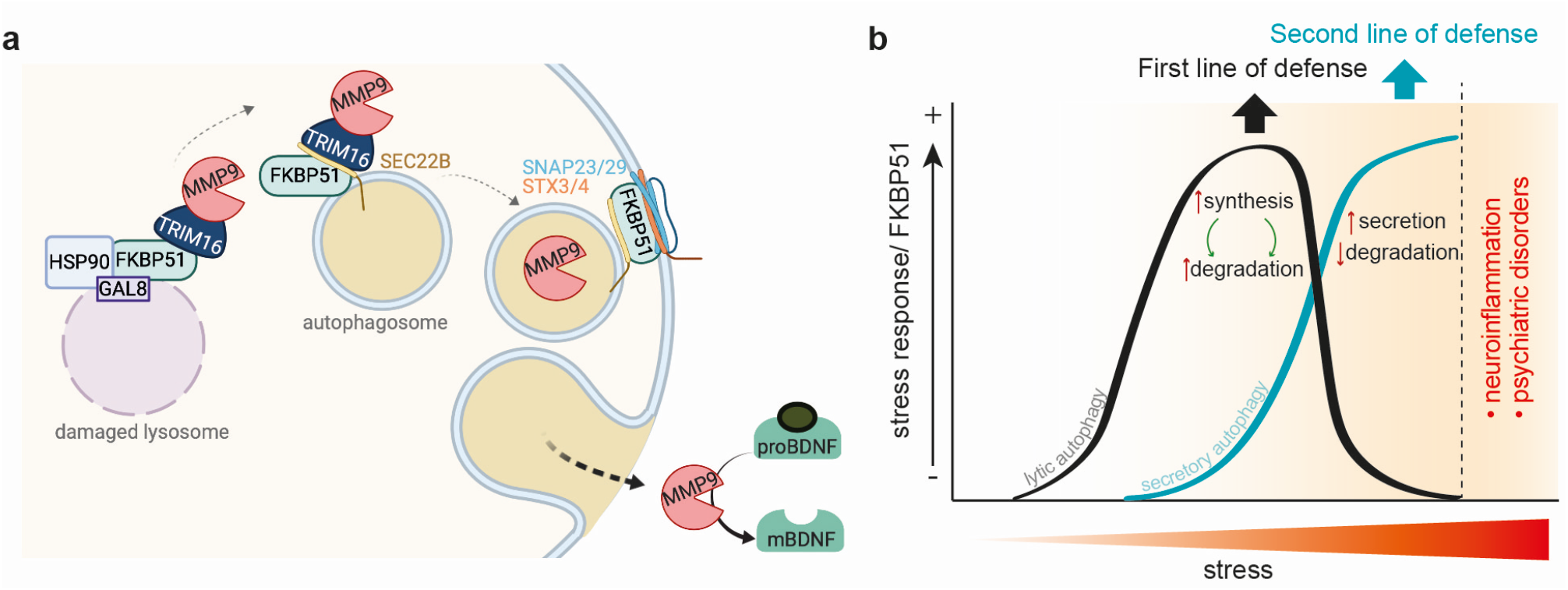
Schematic representation of the findings and proposed model. **a)** Glucocorticoid-mediated stress induces lysosomal damage, which leads to the assembly of FKBP51 with the cargo receptor TRIM16 and the cargo protein (e.g., MMP9). This complex is transported on the autophagosome membrane via the interaction of FKBP51 with SEC22B. The cargo protein is internalized into the autophagosome which is transported to the plasma membrane. The vesicle-plasma membrane fusion is mediated via the SNARE-protein complex assembly, which is regulated by FKBP51 and sensitive to glucocorticoids. The membrane fusion leads to the release of the cargo proteins into the extracellular milieu. The increased secretion of extracellular MMP9 induces the cleavage of proBDNF to its mature form, which becomes the prevailing form in the extracellular space; **b)** A first response to stress triggers lytic autophagy^7^. When the stress persists, a second defense line is activated, which switches the stress response from lytic to secretory autophagy. If stress further persists (e.g., chronic), the initially beneficial proteins secreted in response to stress, might lead to neuroinflammation and psychiatric disorders.

### MMP9 and BDNF – early proteins of the stress response?

Among the identified proteins that are secreted in a stress- and autophagy-dependent manner, MMP9 stands out as key regulator in CNS development and plasticity. As an endopeptidase, MMP9 cleaves-among other substrates – extracellular components, cell adhesion molecules, cell surface receptors, and neurotrophin precursors, such as proBDNF. Hence, MMP9 is implicated in active dendritic spine remodeling and stabilization, pre- and postsynaptic receptor dynamics, and synaptic pruning^27,28^. In our study, we describe secretory autophagy as a novel stress-inducible mechanism for MMP9 secretion both *in vitro* and *in vivo*, elicited respectively by dexamethasone treatment or FS.

Elevated GC concentrations caused by prolonged stress may lead to structural changes and neuronal damage in distinct structures of the brain, which can contribute to the development and progression of neuropsychiatric disorders. Furthermore, accumulating evidence suggests that neuroplasticity, a central mechanism of neuronal adaptation, is altered in stress-related disorders and animal models of stress. Alongside its classical functions as a key neurotrophin during the development of the nervous system, BDNF modulates synaptic plasticity and is involved in stress-induced hippocampal adaptation and pathogenesis of MDD^29^. Our results demonstrate that secretory autophagy-mediated elevation in extracellular MMP9 levels causes an increase in the extracellular mBDNF/proBDNF ratio, a profile that has been observed in MDD patients^30^. MMP9 might, therefore, represent a means to increase the availability of mBDNF, “optimizing” BDNF levels to facilitate synaptic plasticity and remodeling under prolonged stress conditions.

Interestingly, it has been shown that stimulation of primary cortical neurons with BDNF can upregulate MMP9 mRNA and protein levels suggesting a positive feed-forward mechanism for BDNF maturation^31^. In line with these observations, Niculescu *et al*., have elegantly reported that synaptic clustering requires BDNF/TrkB signaling, MMP9 activity and NMDAR activation. While mBDNF/TrkB signaling strengthens clustered synapses, proBDNF/p75^NTR^ signaling depresses out-of-sync synapses via NMDAR activation^32^.

MMP9-mediated conversion of proBDNF to mBDNF occurs at clustered synapses, thereby efficiently sorting synaptic inputs into correlated clusters. Therefore, MMP9/mBDNF/TrkB signaling might be essential for synapses to exhibit plasticity and to modify their function and structure in response to a stressor. This physiological response to stress would allow for increased cognitive vigilance (and possibly resilience to further neuronal insults), making it seem plausible that lowered BDNF levels induce a state of increased vulnerability to stress and MDD. Indeed, numerous studies have described central and peripheral reductions in BDNF levels following chronic stress and in patients with MDD, indicating that optimum levels of BDNF (and its isoforms) are critical to ensure proper neuronal functioning^33^. Of note, having observed a rescuing effect of SAFit1 that could possibly be defined as a stabilizing effect on BDNF, FKBP51 could be a promising target for the prevention of prolonged effects of an acute stress, such as a traumatic event. In fact, BDNF levels tend to normalize in response to several interventions, such as antidepressants^34^ or physical activity^35^. However, despite some consistent findings, other studies report incongruent or contrasting results. Thus, in the light of our study and previous publications, future studies should differentially analyze pro- and mBDNF (instead of total BDNF), which may clarify correlations to physiological and diseased states and improve the characterization of BDNF as a biomarker.

Taken together, our findings show for the first time that stress enhances secretory autophagy leading to increased release of numerous proteins, a large number of which is involved in the immune response. Among these, we found MMP9, whose secretion results in effective cleavage of proBDNF and accumulation of ready to act mBDNF in the extracellular space. These results propose a fast response to stress, which enhances neuroplasticity and cognition, followed however, by a possible increase in neuroinflammation in case of prolonged or chronic stress (model summarized in Fig. 7). The antagonizing effect of SAFit1 in this mechanism is therefore of particular interest as a promising pharmacological tool to contrast psychiatric disorders as well as chronic stress-related neuroinflammation.

## Online Methods

### Cell culture

The human cell lines HEK and SY-SY5Y were cultured at 37 °C, 6% CO2 in Dulbecco’s Modified Eagle Medium (DMEM) high glucose with GlutaMAX (Thermo Fisher, 31331-028), supplemented with 10% fetal bovine serum (Thermo Fisher, 10270-106) and 1% antibiotic-antimycotic (Thermo Fisher, 15240-062).

The murine microglia cell line SIM-A9 was cultured at 37 °C, 6% CO_2_ in Dulbecco’s Modified Eagle Medium (DMEM) high glucose with GlutaMAX (Thermo Fisher, 10566016), supplemented with 10% fetal bovine serum (Thermo Fisher, 10270-106), 5% horse serum (Thermo Fisher, 16050-122) and 1% antibiotic-antimycotic (Thermo Fisher, 15240-062).

### Transfections

With 1x trypsin-EDTA (gibco, 15400-054) detached HEK, SH-SY5Y or SIM-A9 cells (2 ×10^6^) were resuspended in 100 μl of transfection buffer [50 mM Hepes (pH 7.3), 90 mM NaCl, 5 mM KCl, and 0.15 mM CaCl_2_]. Up to 2 μg of plasmid DNA was added to the cell suspension, and electroporation was carried out using the Amaxa 2B-Nucleofector system (Lonza). Cells were replated at a density of 10^5^ cells/cm^2^.

#### Plasmids used

FKBP51-FLAG as described in Wochnik et al. 2005^36^. GFP-SEC22B and Trim16-FLAG were kind gifts by Dr. Vojo Deretic (University of New Mexico).

### CRISPR-Cas9 KO generation

Generation of SIM-A9 *Atg5* KO cell line: using Lipofectamine 3000 transfection reagent (Thermo Fisher Scientific, L3000001), cells were transfected with CRISPR-Cas9 *Atg5* plasmid. 48 hours after transfection 2 μg/ml of puromycin (InvivoGen, ant-pr-1) was added to the medium. After 36 hours the medium was changed, and single clones were manually picked and replated in single wells for expansion. KO clones were selected via western blot analysis.

For the generation of *FKBP5* KO SH-SY5Y cell line, cells were co-transfected with a pool of three CRISPR/Cas9 plasmids containing gRNA targeting human *FKBP5* and a GFP reporter (Santa Cruz, sc-401560) together with a homology directed repair plasmid (sc-401560-HDR) consisting of 3 plasmids, each containing a homology-directed DNA repair (HDR) template corresponding to the cut sites generated by the FKBP51 CRISPR/Cas9 KO Plasmid (sc-401560). 48 hours after transfection, medium was changed to 2 μg/ml puromycin-containing medium. After 36 hours single clones were manually picked and replated in single wells for expansion. KO clones were selected via western blot analyses.

#### Plasmids used

FKBP51 CRISPR/ Cas9 KO Plasmid (h), Santa Cruz #sc-401560. pSpCas9 BB-2A-Puro (PX459). V2.0 vector containing Atg5-targeting gRNA (Atg5 [house mouse] CRISPR gRNA 2: TATCCCCTTTAGAATATATC) purchased from GenScript.

### Western Blot Analysis

Protein extracts were obtained by lysing cells in 62.5 mM Tris, 2% SDS, and 10% sucrose, supplemented with protease (Sigma, P2714) and phosphatase (Roche, 04906837001) inhibitor cocktails. Samples were sonicated and heated at 95 °C for 10 min. Proteins were separated by SDS-PAGE and electro-transferred onto nitrocellulose membranes. Blots were placed in Tris-buffered saline solution supplemented with 0.05% Tween (Sigma, P2287) (TBS-T) and 5% non-fat milk for 1 hour at room temperature and then incubated with primary antibody (diluted in TBS-T) overnight at 4 °C. Subsequently, blots were washed and probed with the respective horseradish-peroxidase-or fluorophore-conjugated secondary antibody for 1 hour at room temperature. The immuno-reactive bands were visualized either using ECL detection reagent (Millipore, WBKL0500) or directly by excitation of the respective fluorophore. Recording of the band intensities was performed with the ChemiDoc MP system from Bio-Rad.

#### Quantification

All protein data were normalized to ACTIN or GAPDH, which was detected on the same blot. In the case of LC3B the direct ration of LC3BII over LC3BI is provided.

#### Primary antibodies used

LC3B-II/I (1:1000, Cell Signaling, #2775), FLAG (1:7000, Rockland, 600-401-383), FKBP51 (1:1000, Bethyl, A301-430A), ACTIN (1:5000, Santa Cruz Biotechnology, sc-1616), GAPDH (1:8000, Millipore CB1001), TRIM16 (1:1’000, Bethyl A301-160A), CTSD (for Mouse) (1:50, Abcam, ABCAAB6313-100), SNAP29 (1:1000, Sigma, SAB1408650), SNAP23 (1:1000, Sigma, SAB2102251), STX3 (1:1000, Sigma, SAB2701366), STX4 (1:1000, Cell Signalling. #2400), GAL8 (1:1’000, Santa Cruz, sc-28254), GAL3 (1:1’000, Santa Cruz, sc-32790), SEC22B (1:1000, Abcam, ab181076)

### Co-immunoprecipitation

In case of FLAG-or GFP-tag immunoprecipitation, cells were cultured for 3 days after transfection. Cells were lysed in CoIP buffer [20 mM tris-HCl (pH 8.0), 100 mM NaCl, 1 mM EDTA, and 0.5% Igepal complemented with protease (Sigma) and phosphatase (Roche, 04906837001) inhibitor cocktail] for 20 min at 4 °C with constant mixing. The lysates were cleared by centrifugation, and the protein concentration was determined and adjusted (1.2 mg/ml); 1 ml of lysate was incubated with 2.5 μg of FLAG or GFP antibody overnight at 4 °C with constant rotating. Subsequently, 20 μl of bovine serum albumin– blocked protein G Dynabeads (Invitrogen, 100-03D) were added to the lysate-antibody mix followed by a 3 hour incubation at 4 °C. Beads were washed three times with PBS, and bound proteins were eluted with 100 μl of 1 × FLAG peptide solution (100 to 200 ug/ml, Sigma F3290) in PBS for 30 min at 4 °C. In case of precipitation of GFP tag, elution was performed by adding 60 μl of Laemmli sample buffer and by incubation at 95 °C for 5 min. 5 to 15 μg of the input lysates or 2.5 to 5 μl of the immunoprecipitates were separated by SDS–polyacrylamide gel electrophoresis (PAGE) and analyzed by western blotting. When quantifying co-immunoprecipitated proteins, their signals were normalized to input protein and to the precipitated interactor protein.

### Tandem fluorescent-tagged Galectin-3 (tfGal3) assay

SH-SY5Y cells were transfected with a the ptfGalectin3 plasmid (Addgene.org, #64149) that expresses Galectin3 tagged with both EGFP (inactivated in lysosomes) and mRFP (resists inactivation in lysosomes)^16^. Transfection in SH-SY5Y cells was performed using Lipofectamine 3000 transfection reagent (Thermo Fisher Scientific, L3000001), followed by drug treatment the next day. Cells were fixed (1% PFA for 1 h) one day later and analyzed by laser scanning microscopy (Leica Confocal Sp8). Vesicles were counted by an experimenter blind to the conditions.

### Quantitative PCR (RT-qPCR) Analysis

Total RNA was isolated from SIMA-A9 cells using the RNeasy Mini Kit (Qiagen, 74104). 5 μg of total RNA were reverse transcribed using a High Capacity cDNA Reverse Transcription Kit (Thermo Fisher, 4368814). Quantitative PCRs were performed using the TaqMan StepOnePlus Real-Time PCR System and a TaqMan 5′-nuclease probe method (Thermo Fisher). All transcripts were normalized to *Hprt* and *Gapdh*. Predesigned human TaqMan assays (Thermo Fisher) were used for quantifying gene expression of *Bdnf* (Mm04230607_s1), *Hprt* (Mm03024075_m1), and *Gapdh* (Mm99999915_g1).

### ELISA

The solid-phase sandwich ELISA (enzyme-linked immunosorbent assay) for the following antibody detection was performed according to the manufacturer protocol: CTSD (abcam ab213845), MMP9 (abnova KA0398), proBDNF (AssaySolutions AYQ-E10246) or mBDNF (AssaySolutions AYQ-E10225). Briefly, microwells were coated with mouse antibody followed by a first incubation with biotin-coupled anti mouse antibody, a second incubation with streptavidin-HRP and a final incubation with the SIM-A9 culture medium. Amounts of respective proteins were detected with a plate reader (iMARK, Bio-Rad) at 450 nm.

### Animal husbandry and ethics

C57BL/6 male mice (Martinsried, Germany), FKBP51-KO and respective WT males (Martinsried, Germany) at 8–12 weeks of age were housed in cages of 4-5 at 21 °C with a 12:12 h light-dark cycle with food and water ad libitum before experiments. Allocation of animals to experimental groups in respect to date of birth and litter was done using random number generation approach (RAN function in Excel). All procedures were done in accordance with European Communities Council Directive 2010/63/EU and approved by Government of Upper Bavaria.

### *In vivo* brain microdialysis in mice

#### Surgeries

Surgeries were performed as described previously in Anderzhanova et al., 2013^37^ under 2 % isoflurane in oxygen (Abbot, India), Metacam® 0.5 mg/ kg, s.c (Boehringer Ingelheim GmbH, Germany), and Novalgine 200 mg/kg, s.c. (Sanofi-Aventis, Germany) systemic anesthesia and analgesia. Lidocaine 2% (Xylocaine, AstraZeneca) was used for local anesthesia. Coordinates for microdialysis probe guide cannula implantations into the right mPFC were set in accordance to Paxinos and Franklin Mouse Brain Atlas^38^ with bregma as a reference point: AP 2.00 mm, ML 0.35 mm, and DV −1.50 mm. Stereotaxic manipulations were performed in the TSE stereotaxic frame (TSE Systems Inc., Germany). Animals were allowed for 7 days of recovery in individual microdialysis cages (16 × 16 × 32 cm3). Metacam (0.5 mg/kg, s.c) was injected within the first three days after surgeries, when required.

#### Microdialysis

The perfusion setup was the line comprised of FET tubing of 0.15 mm ID (Microbiotech Se, Sweden), a 15 cm-PVC inset tubing (0.19 mm ID), a dual-channel liquid swivel (Microbiotech Se, Sweden). Perfusion medium was sterile RNase free Ringer’s solution (BooScientific, USA) containing 1% bovine serum albumin (BSA) (Sigma-Aldrich, Cat.N A9418). Perfusion medium was delivered to the probe at the flow rate of 0.38 μl/ min with the syringe pump (Harvard Apparatus, USA) and withdrawn with the peristaltic pump MP2 (Elemental Scientific, USA) at the flow rate of 0.4 μl/ min. Microdialysis CMA 12 HighCO Metal Free Probe was of 2 mm length membrane with 100 kDa cut off (Cat.N. 8011222, CMA Microdialysis, Sweden). All lines were treated with 5 % polyethylenimine (PEI) for 16 hrs and then with H_2_O for 24 hrs before experiments. The microdialysis probe was inserted into the implanted guide cannula (under 1-1.5 min isoflurane anesthesia, 2% in air) 6 days after the stereotaxic surgery and 18 h before the samples collection. A baseline sample collection phase (three samples) was always preceding the FS, which allowed us to express the changes in the extracellular content of proteins as relative to the baseline values. On the experimental day, microdialysis fractions were constantly collected (for 30 min) into Protein LoBind tubes (Eppendorf, Germany) at a perfusion rate of 0.4 μl/min. During collection time, tubes were kept on ice. After collection of three baseline samples animals were transferred to the foot shock (FS) chamber (ENV-407, ENV-307A; MED Associates, 7 St Albans, VT, USA) connected to constant electric flow generator (ENV-414; MED Associates) and a FS (1.5 mA × 1 sec × 2) was delivered. After this procedure, mice were returned to the microdialysis cage where two post-FS samples were collected. To examine an effect of ULK1 inhibitor MRT 68921 on stress-evoked changes in extracellular content of proteins, the drug was injected intraperitoneally in a dose of 5.0 mg/kg and in a volume 10 ml/kg four hours before the moment of FS (the drug was prepared freshly dissolving a stock solution [60%EthOH/40% DMSO mixture] with saline in proportion 1:20). 30 min microdialysis fractions were collected on ice into 1.5 ml protein LoBind tubes (Eppendorf, Germany) preloaded with 0.5 μl protease inhibitor cocktail 1:50 (Roche) and then immediately frozen on dry ice. At the end of the experiment, probes were removed, brains were frozen and kept at −80 °C for the probe placement verification. 40 μm brain sections were stained with cresyl violet (Carl Roth GmbH, Germany) and probe placement was verified under a microscope using Paxinos and Franklin (2001) mouse atlas^38^. If probe placement was found to be out of the targeted region of interest, the respective samples were excluded from the study.

### Interactome analyses

#### Sample preparation

All samples were prepared in quadruplicate. HEK cells were transfected with a FLAG tagged FKBP51 expressing plasmid or a FLAG expressing control vector. 48 hours after transfection, a FLAG IP was performed on all the samples and the eluted proteins were separated by SDS gel electrophoresis. Separated proteins were stained with Coomassie staining solution for 20 min and destained over night with Coomassie destaining solution. Each gel lane was cut into 21 approximately 2.5 mm slices per biological replicate and these further cut into smaller gel pieces.

#### In-gel-trypsin digestion and peptide extraction

The gel pieces were covered with 100μl of 25 mM NH_4_HCO_3_/50% acetonitrile in order to destain the gel pieces completely and mixed for 10 min at room temperature. The supernatant was discarded, and this step was repeated twice. Proteins inside the gel pieces are reduced with 75μl of100 mM dithiothreitol/25 mM NH_4_HCO_3_ mixed at 56 °C for 30 min in the dark. The supernatant was discarded and 100μl of 200 mM iodoacetamide were added for alkylation to the gel pieces and mixed for 30 min at room temperature. The supernatant was discarded, and the gel pieces washed twice with 100μl 25 mMNa_4_HCO_3_/50% acetonitrile and mixed for 10 min at room temperature. The supernatant was discarded, and gel pieces were dried for approximately 20 min at room temperature. Proteins were digested with 50 μl trypsin solution [5 ng/μl trypsin/25 mM NH_4_HCO_3_] over night at 37 °C. Peptides were extracted from the gel pieces by mixing them in 50 μl of 2% formic acid/50% acetonitrile for 20 min at 37 °C and subsequently sonicating them for 5 min. This step was repeated twice with 50 μl of 1% formic acid/50% acetonitrile. The supernatants of three slides were pooled. Each sample generated 7 samples to be submitted to LC-MS/MS analysis.

#### LC MS/MS

Tryptic peptides were then dissolved in 0.1% formic acid and analyzed with a nanoflow HPLC-2D system (Eksigent, Dublin, CA, USA) coupled online to an LTQ-Orbitrap XL™ mass spectrometer. Samples were on-line desalted for 10 min with 0.1% formic acid at 3 μl/min (Zorbax-C18 (5 μm) guard column, 300 μm × 5 mm; Agilent Technologies, Santa Clara, CA, USA) and separated via RP-C18 (Dr. Maisch, Germany, 3 μm) chromatography (in-house packed Pico-frit column, 75 μm × 15 cm, New Objective, Woburn, MA, USA). Peptides were eluted with a gradient of 95% acetonitrile/0.1% formic acid from 10% to 45% over 93 min at a flow rate of 200 nl/min. Column effluents were directly infused into the mass spectrometer via a nano-electrospray ion source (Thermo Fisher Scientific). The mass spectrometer was operated in positive mode applying a data-dependent scan switch between MS and MS/MS acquisition. Full scans were recorded in the Orbitrap mass analyzer (profile mode, m/z 380–1600, resolution R = 60000 at m/z 400). The MS/MS analyses of the five most intense peptide ions for each scan were recorded in the LTQ mass analyzer in centroid mode.

#### Peptides and proteins identification

The MS raw data were first converted in a peaks list using Bioworks v. 3.3.1 (Thermo Fischer Scientific, San Jose, CA) and then were searched against the SwissProt_15.3 (uniprot 29.05.09) human database using Mascot search algorithm (v. 2.2.07, www.matrix.com)

The precursor and fragment mass tolerance were set to 20 ppm and 0.8 Da, respectively. Trypsin/P including one missed cleavage was set as enzyme. Methionine oxidation and carbamidomethylation of cysteine were searched as variable and static modifications, respectively. The significance level in protein identification was set p>0.95, the peptide score cut-off was set 20. Proteins identified with at least 3 peptides were considered for the protein data processing.

The identified proteins overlapping in the four replicates were considered as protein interactors.

The mass spectrometry proteomics data have been deposited to the ProteomeXchange Consortium via the PRIDE^39^ partner repository with the dataset identifier PXD017328 and 10.6019/PXD017328

### Secretome analyses

#### Sample preparation

All samples were prepared in triplicates. For each replicate 2 × 150 mm culture dishes, containing each 20 ml of medium, were used. WT and *Atg5* KO SIM-A9 cells were cultured in methionine-free medium [DMEM, high glucose, no glutamine, no methionine, no cystine (Thermo Fisher, 21013024), 2 mM L-Glutamine (Thermo Fisher, 25030081), 1 mM sodium pyruvate (Thermo Fisher, 11360070), 0.2 mM L-Cysteine (Sigma, C7352), 10% dialyzed fetal bovine serum (Thermo Fisher, 30067334), 1% Antibiotic-Antimycotic (Thermo Fisher, 15240-062)] for 30 min (12 hours and 30 min before supernatant collection). After 30 min, medium was changed to AHA enriched medium [methionine-free medium with 20 μM Click-IT™ AHA (L-Azidohomoalanine) (Thermo Fisher, C10102)] to label newly synthesized proteins. After 6 hours, cells were treated with 100 nM dexamethasone or DMSO (vehicle) at a dilution of 1:10’000 (6 hours before supernatant collection). After 3 hours (3 hours before supernatant collection), all media were refreshed in order to analyze proteins that were secreted only during the last 3-hours time-window. After 3 hours, supernatants were collected and centrifuged at 1000 × g for 5 min. EDTA-free proteinase inhibitor cocktail (Sigma, S8830) was added and supernatants were frozen at −20°C. The next day, supernatants were thawed on ice and concentrated with Amicon Ultra 15 centrifugal filters, Ultracell 3K (Millipore, UFC900324) at 4000 × g at 4 °C for 5 hours. The concentrated samples were transferred into new 1.5 ml tubes (Sigma, Z606340). For the enrichment and digestion of AHA-labeled proteins, the Click-iT™ Protein Enrichment Kit (Thermo Fisher Scientific) was used according to the instructions of the supplier, despite that only half of the suggested volumes were used^18^. Resulting peptides were further desalted by using 0.1 % formic acid in 50 % acetonitrile in the micro-column format (three discs, Ø 1.5 mm, C18 material, 3M Empore per micro-column were used)^40^. After drying in a centrifugal evaporator, the samples were stored at −20 °C until LC-MS/MS analysis.

#### LC-MS/MS

LC-MS/MS measurement of peptides in eluates was performed using a nanoLC UltiMate 3000 (Thermo Fisher Scientific) coupled to a quadrupole-Orbitrap Q Exactive HF-X mass spectrometer (Thermo Fisher Scientific). Peptides separated on an Acclaim PepMap analytical column (0.1 mm × 15 cm, C18, 2 μM, 100 Å; Thermo Fisher Scientific) using a 60 min linear gradient from 3-28 % solvent B [0.1 % formic acid, 5 % DMSO in acetonitrile] in solvent A [0. 1% formic acid, 5 % DMSO in water] at a flow rate of 10 μL/min. The mass spectrometer was operated in data dependent acquisition and positive ionization mode. MS1 spectra were acquired over a range of 360-1300 m/z at a resolution of 60’000 in the Orbitrap by applying an automatic gain control (AGC) of 3e6 or maximum injection time of 50 ms. Up to 12 peptide precursors were selected for fragmentation by higher energy collision-induced dissociation (HCD; 1.3 m/z isolation window, AGC value of 1e5, maximum injection time of 22 ms) using 28% normalized collision energy (NCE) and analyzed at a resolution of 15’000 in the Orbitrap.

#### Peptide and Protein identification and quantification

Peptide and protein identification and quantification was performed using MaxQuant (version 1.6.0.16)^41^ by searching the tandem MS data against all murine canonical and isoform protein sequences as annotated in the Swissprot reference database (25175 entries, downloaded 13.07.2018) using the embedded search engine Andromeda^42^. Carbamidomethylated cysteine was set as fixed modification and oxidation of methionine and N-terminal protein acetylation as variable modification. Trypsin/P was specified as the proteolytic enzyme and up to two missed cleavage sites were allowed. Precursor tolerance was set to 4.5 ppm and fragment ion tolerance to 20 ppm. The minimum peptide length was set to seven and all data were adjusted to 1% PSM and 1 % protein FDR. Intensity-based absolute quantification (iBAQ)^43^ was enabled within MaxQuant.

#### Data analysis

The Perseus software suite (v. 1.6.2.3)^44^ was used to filter out contaminants, reverse hits and protein groups, which were only identified by site. Only protein groups that were detected in at least two out of the three replicates in at least one condition were considered for the analysis.

The filtered data was log normalized and missing values were imputed according to the normal distributed imputation algorithm implemented in the Perseus framework. Default values were used (width: 0.3; down shift: 1.8). To find the significantly regulated protein groups a multiple t-test (volcano plot) analysis was performed with a difference of s0 =1 and a false discovery rate (FDR) of 0.01.

Proteins with a significant (FDR<0.01) and at least 2-fold (s0>1) increased secretion in the WT compared to the Atg5 KO samples were analyzed with the reactome pathway browser (www.reactome.org; pathway browser version 3.6, reactome database release 70)^45^. The same list was also analyzed with the SynGO knowledgebase (www.syngoportal.org)^19^.

#### Data availability

The mass spectrometry proteomics data have been deposited to the ProteomeXchange Consortium via the PRIDE partner repository with the dataset identifier PXD017076. Reviewer account details: Username. Password: GT6DRDWo.

### Automated literature search

Protein lists resulting from the interactome and secretome analyses were subjected to an automated literature search performed with a custom python code, revealing as output the number of references found for each search. For the interactome list, the algorithm performed an automated PubMed search of the protein with “autophagy”, “proteostasis”, or “ubiquitin proteasome system”. For the secretome list, the algorithm performed an automated PubMed search of each protein with “neuroplasticity”. False negatives resulting from protein name or abbreviation ambiguity were manually corrected.

#### Code for interactome analysis

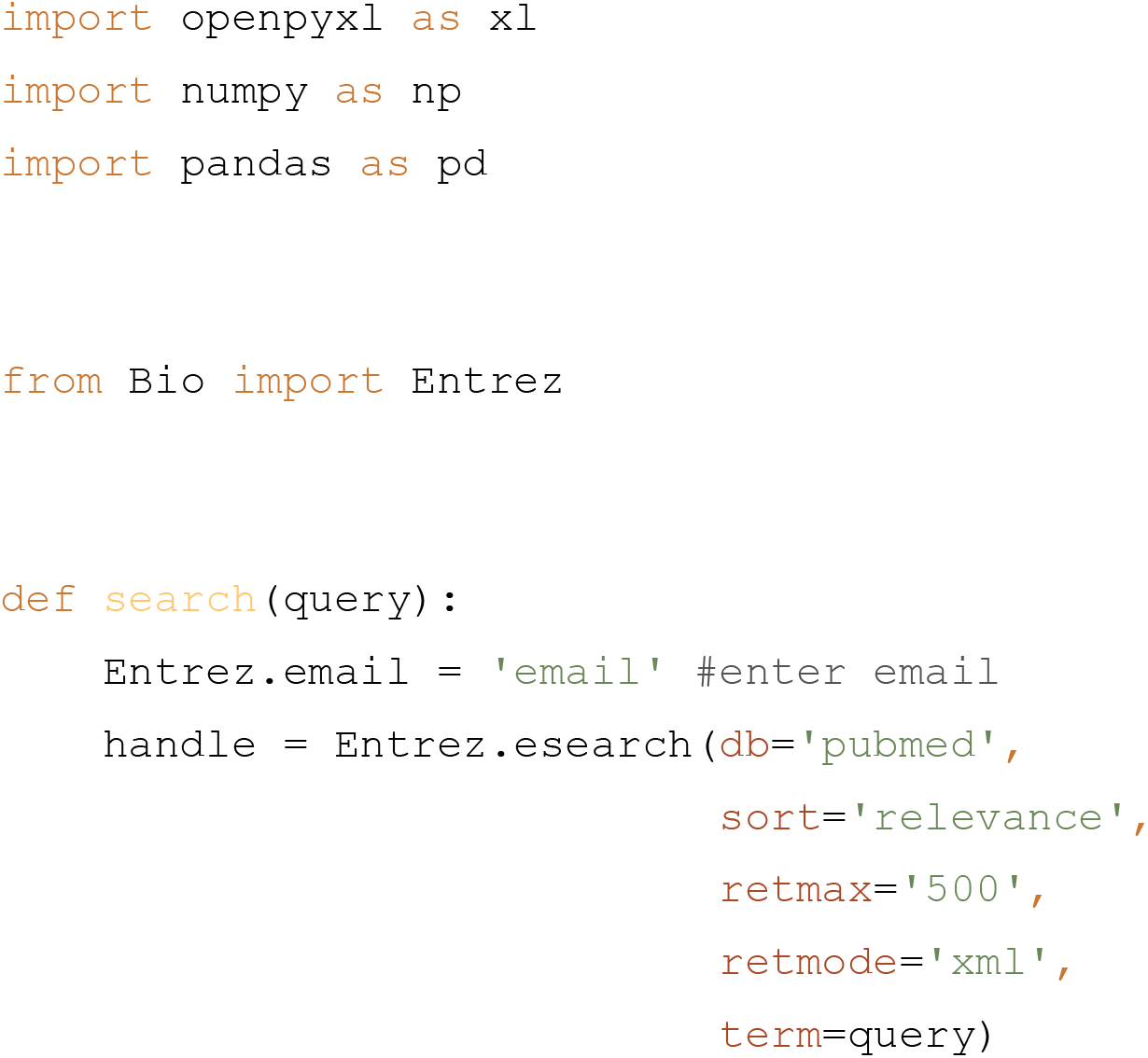
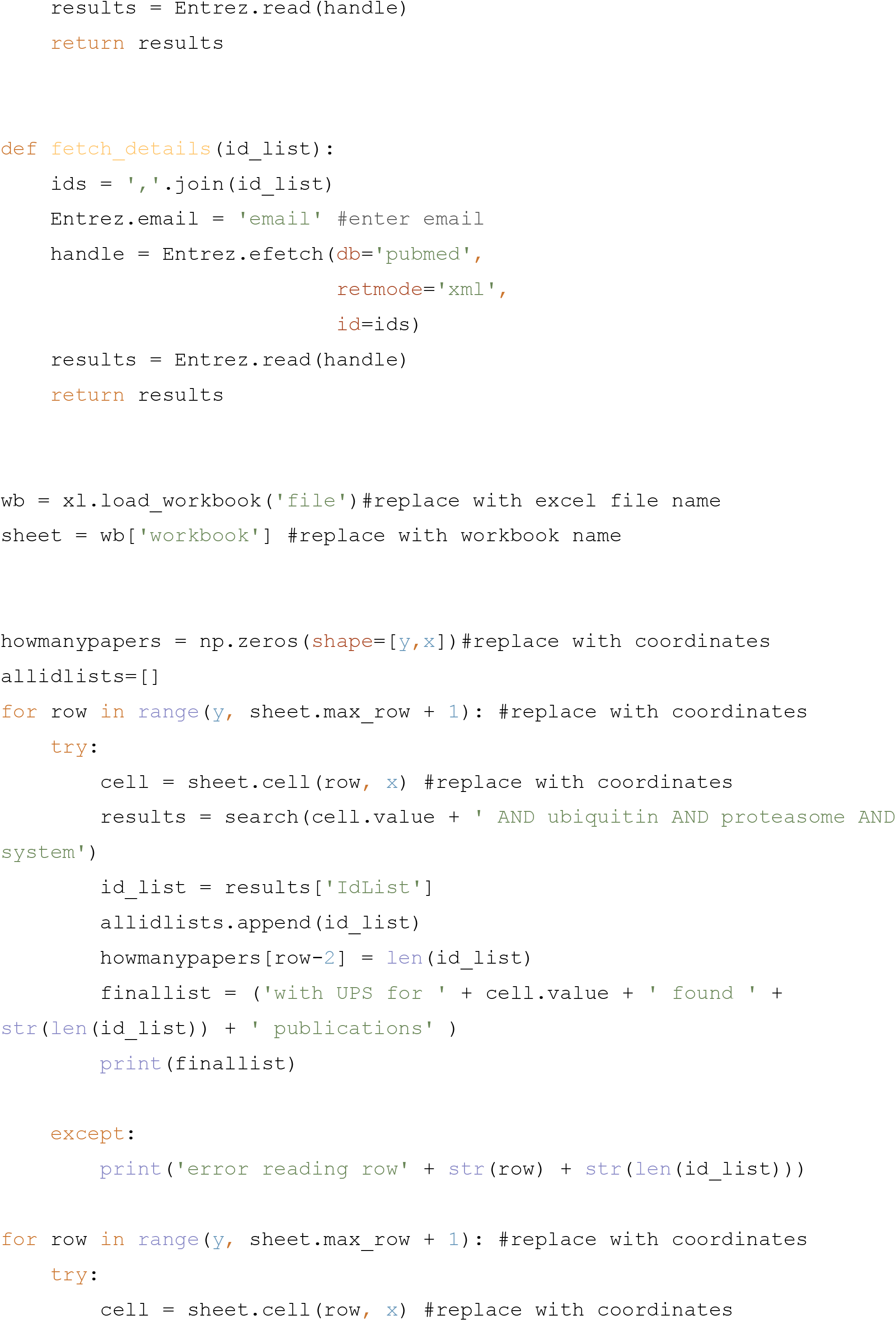
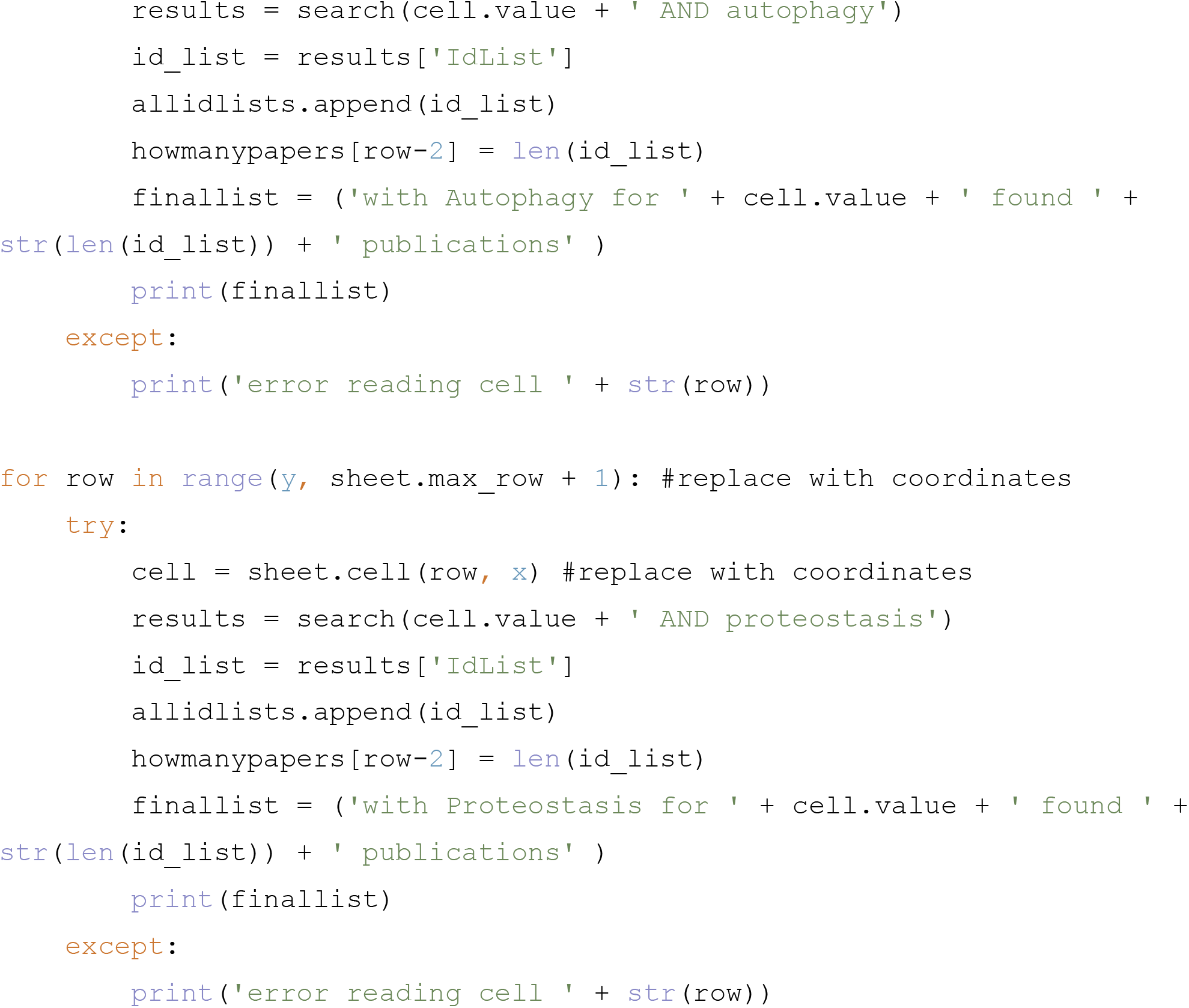

#### Code for secretome analysis

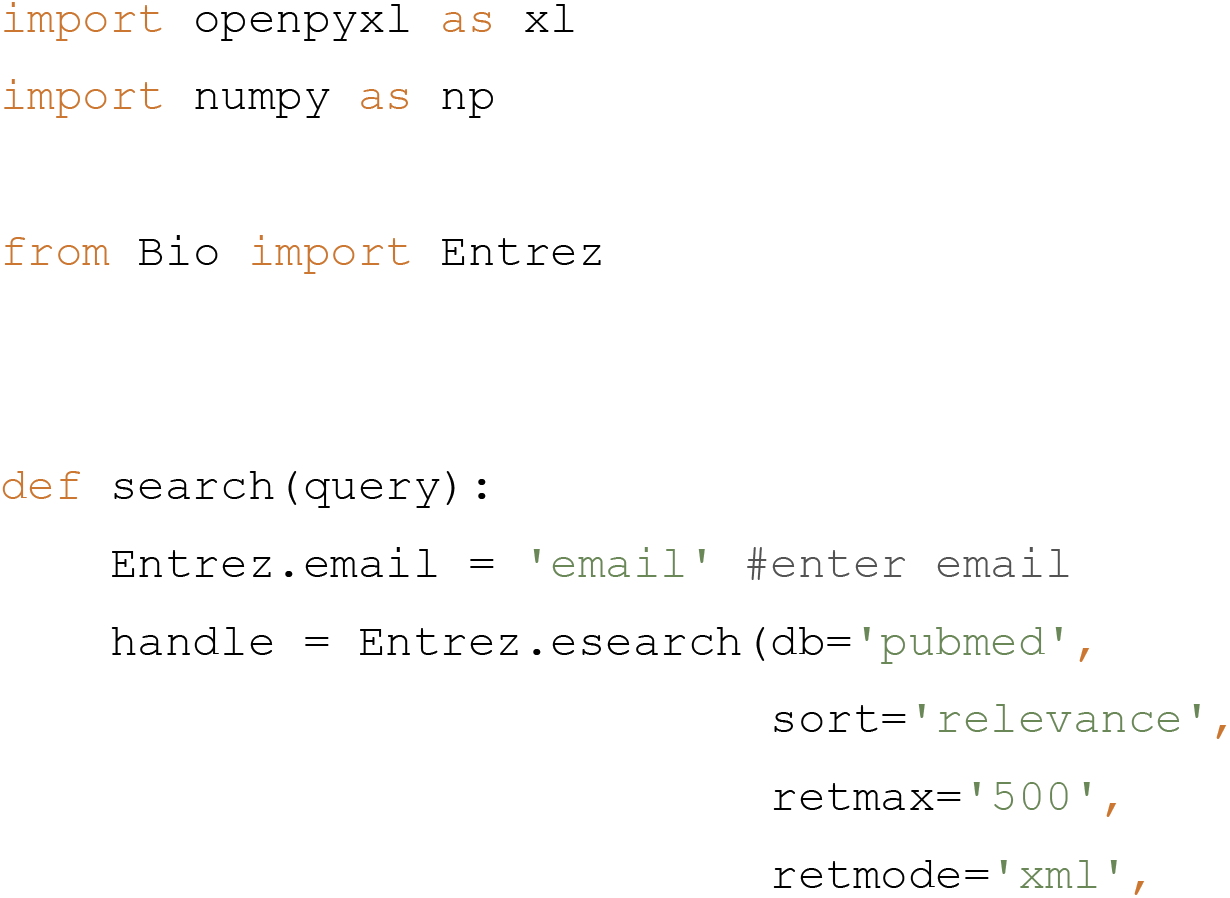
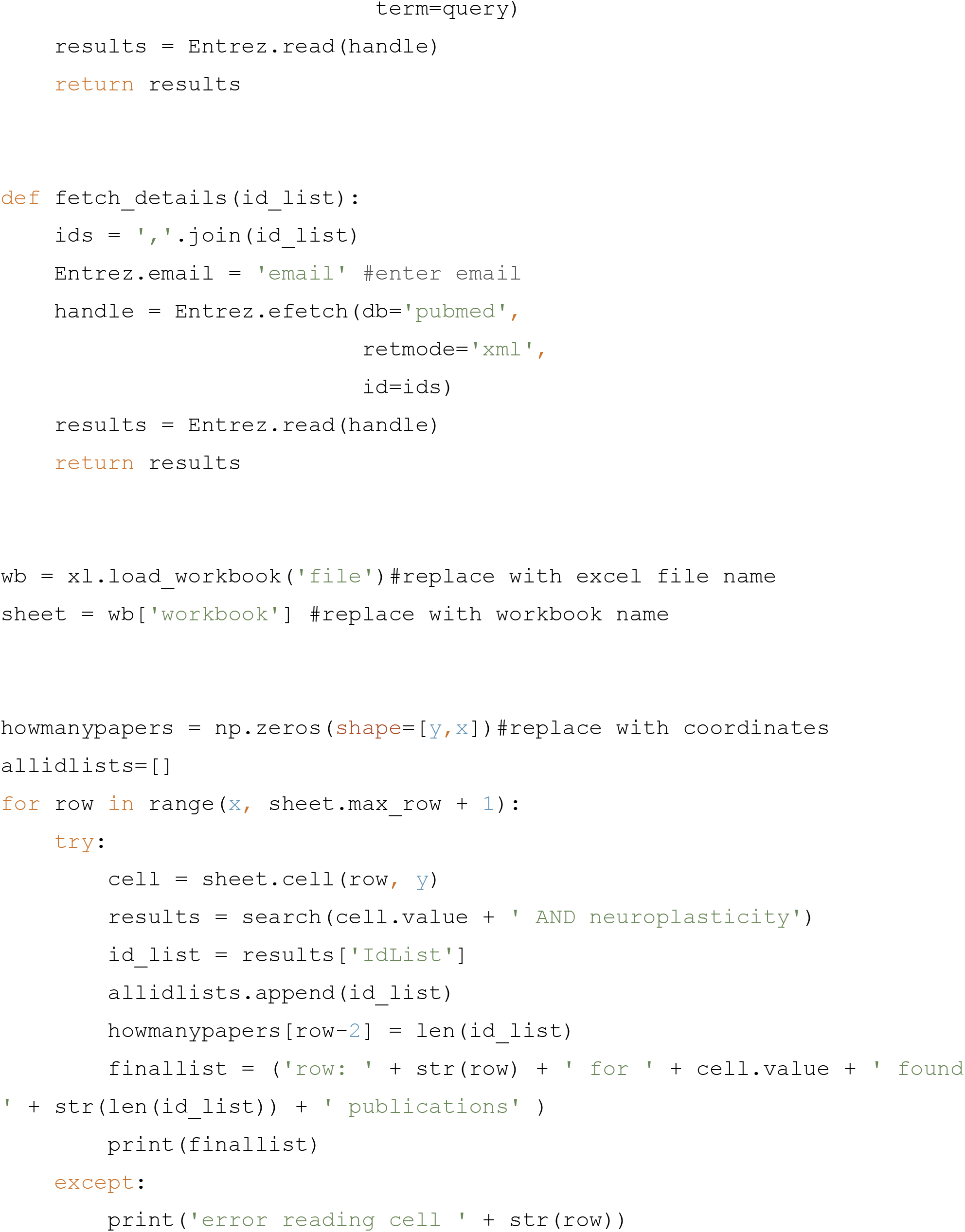

### Statistical analyses

Except from proteomics data, all statistical analyses were performed with Prism version 8.1.1 (GraphPad Software, La Jolla California USA, www.graphpad.com). Data distribution was tested with the Shapiro-Wilk test. Individual data points are shown when possible. Error bars indicate standard error mean (s.e.m.). Analyses of paired measurements were performed using the Sidak’s multiple comparisons test for individual comparisons and two-way ANOVA test for group comparisons. Unpaired analyses were performed using one-way unpaired t-test with appropriate correction depending on the distribution and variance: normally distributed data with different variance were corrected with Welch’s correction; non-normally distributed data were analyzed with Mann Whitney test. For multiple comparisons of normally distributed data, Tukey’s comparison test was applied, while for non-normally distributed data, the Kruskal Wallis test was performed. P < 0.05 was considered statistically significant.

## Acknowledgements

Many thanks to Elisabeth Binder for her support and help to this study. This study was funded by a NARSAD Young Investigator Award by Brain and Behavior Research Foundation, honored by P&S Fund (Awarded to Nils C Gassen, Grant ID 25348).

## Author Contributions

SM, EAA and NCG designed the study and performed the experiments. SM and NCG analyzed the data and wrote the manuscript. SW, FD, KW and GM performed and analyzed the proteomic experiments. KH, TB, JH, MLP and LJ contributed to the experiments. NCG supervised the study. All authors contributed to the final version of the manuscript.

