## Supplementary information for "Stress-primed secretory autophagy drives extracellular BDNF maturation"

### Supplementary Figures

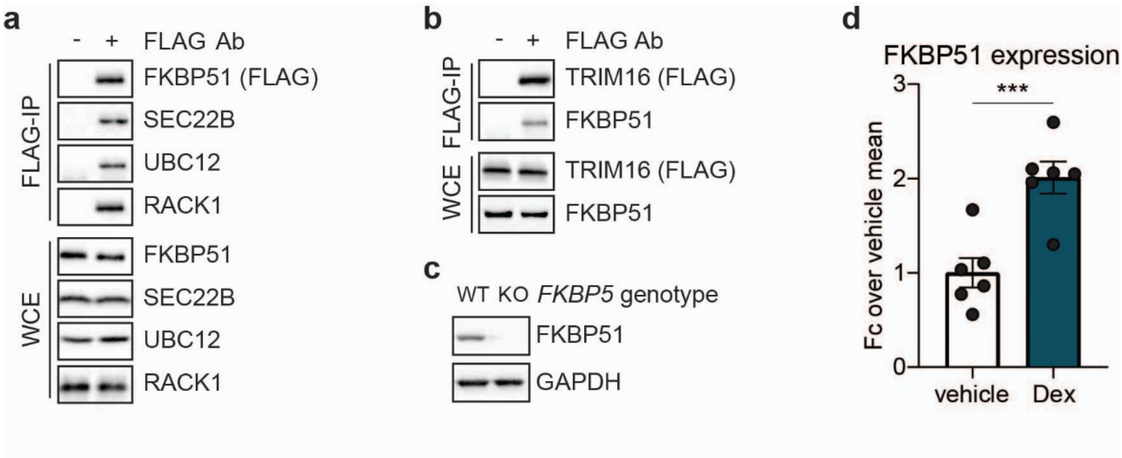


**Figure S1: Control western blots relative to Figure 1**

**a)** Western blotting for FKBP51, UBC12 and RACK1 in FLAG-tagged FKBP51 co-IP (FLAG-IP) and whole cell extract (WCE) as control. **b)** Western blotting for TRIM16 and FKBP51 in FLAG-tagged TRIM16 co-IP (FLAG-IP) and whole cell extract (WCE) as control. **c)** Western blotting for FKBP51 in WT and *FKBP5* KO SH-SY5Y cells. **d)** Quantifications of western blots for FKBP51 in SH-SY5Y cells treated with vehicle or 100 nM dexamethasone (Dex) for 4 hours. $n = 6$. Unpaired one-tailed t-test was performed; ***$P < 0.001$. Data shown as mean ± s.e.m. Ab, antibody; Fc, fold change.

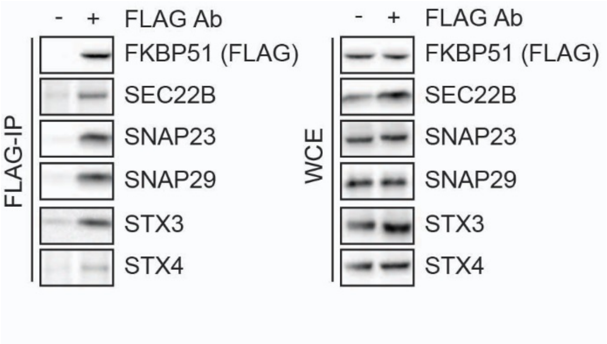


**Figure S2: Control co-IP in SIM-A9 cells**

Western blotting for FKBP51, SEC22B, SNAP23, SNAP29, STX3 and STX4 in FLAG-tagged FKBP51 co-IP (FLAG-IP) performed in SIM-A9 cells and whole cell extract (WCE) as control. Ab, antibody.

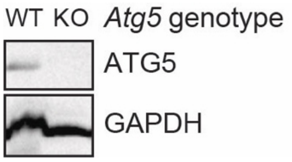


**Figure S3: KO validation western blot**

Western blotting for ATG5 in WT and *Atg5* KO SIM-A9 cells.

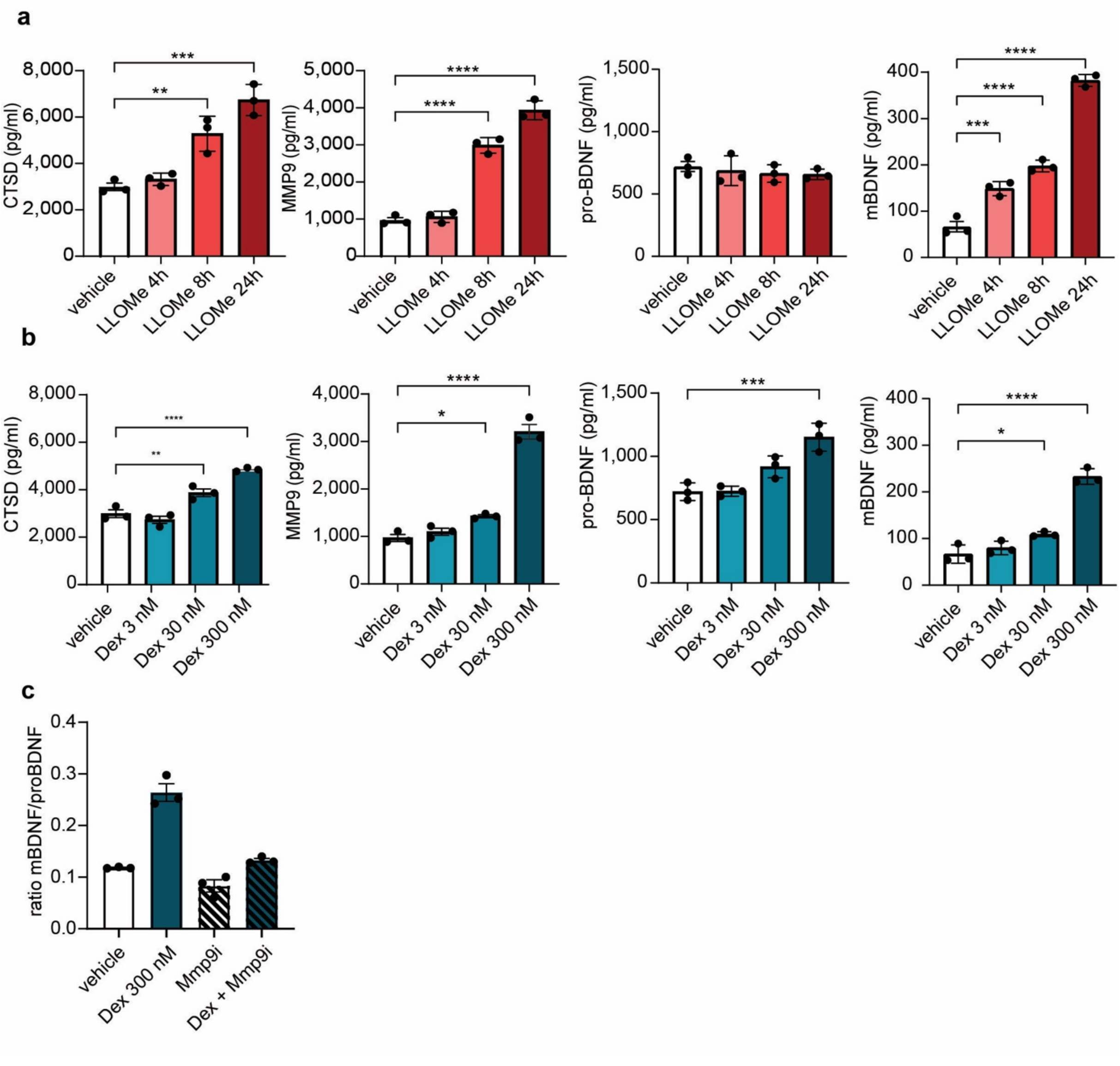


**Figure S4: ELISAs of SIM-A 9 supernatants**

**a)** CTSD, MMP9**,** proBDNF, mBDNF levels from supernatants measured via ELISA after WT SIM-A9 cells were treated with 1 mM LLOMe for 4, 8 or 24 hours or vehicle or 24 hours; **b)** CTSD, MMP9**,** proBDNF, mBDNF levels from supernatants measured via ELISA after WT SIM-A9 cells were treated with 3 nM, 30 nM or 300 nM dexamethasone (Dex) or vehicle for 4 hours; **c)** mBDNF/proBDNF ratio from supernatants measured via ELISA after WT SIM-A9 cells were treated with 300 nM Dex, Dex + Mmp9 inhibitor I (MMP9i), or vehicle for 4 hours. Tukey's multiple comparison tests were performed; *P < 0.05; **P < 0.01; ***P < 0.001; ****P < 0.0001. Data shown as mean ± s.e.m.

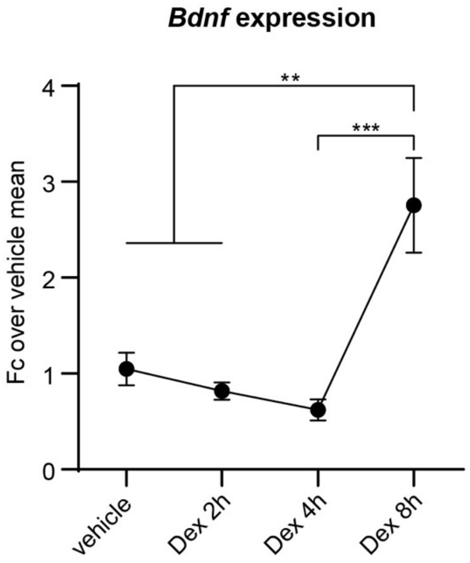


**Figure S5: *Bdnf* expression in SIM-A9 cells**

Quantitative polymerase chain reaction (qPCR) of *Bdnf* expressed in SIM-A9 cells treated with vehicle or 100 nM Dex for 2, 4 and 8 hours. Tukey's multiple comparison test was performed; **P < 0.01; ***P < 0.001. Data shown as mean ± s.e.m.

### Supplementary references

### List of references to neuroplasticity-related proteins listed in Table 1

| Protein | Reference |
|---|---|
| APP | 1–3 |
| JUN | 4,5 |
| VIM | 6,7 |
| CDC42 | 8,9 |
| MMP9 | 10–12 |
| CCL2 | 13,14 |
| EEF2 | 15–17 |
| HMGB1 | 18 |
| IGF1 | 19–21 |
| KDR | 22,23 |
| LRP1 | 24–27 |
| PAK1 | 28,29 |
| PAK3 | 30,31 |
| STMN2 | 32,33 |
| TF | 34 |

| | |
|---|---|
| TGFB1 | 35–37 |
| CAT | 38,39 |
| HMOX1 | 40 |
| JUNB | 41 |

37. Jaskova, K., Pavlovicova, M., Cagalinec, M., Lacinova, L. & Jurkovicova, D. TGFβ1

downregulates neurite outgrowth, expression of Ca2+ transporters, and mitochondrial dynamics of in vitro cerebellar granule cells. *Neuroreport* **25**, 340–346 (2014).
